# Nuclear Abl Drives miR-34c Transfer by Extracellular Vesicles to Induce Radiation Bystander Effects

**DOI:** 10.1101/209767

**Authors:** Shubhra Rastogi, Amini Hwang, Josolyn Chan, Jean YJ Wang

## Abstract

**SUMMARY:** Ionizing radiation stimulates nuclear accumulation of Abl tyrosine kinase that is required for directly irradiated cells to produce microRNA-34c-containing extracellular vesicles, which transfer the microRNA into non-irradiated cells to induce reactive oxygen species and bystander DNA damage.

**ABSTRACT:** Ionizing radiation (IR) activates an array of DNA damage response (DDR) that includes the induction of bystander effects (BE) in cells not targeted by radiation. How DDR pathways in irradiated cells stimulate BE in non-targeted cells is mostly unknown. We show here that extracellular vesicles from irradiated cells (EV-IR) induce reactive oxygen species (ROS) and DNA damage when internalized by un-irradiated cells. We found that EV-IR from Abl-NLS-mutated cells could not induce ROS or DNA damage, and restoration of nuclear Abl rescued those defects. Expanding a previous finding that Abl stimulates miR-34c expression, we show here that nuclear Abl also drives the vesicular secretion of miR-34c. Ectopic miR-34c expression, without irradiation, generated EV-miR-34c capable of inducing ROS and DNA damage. Furthermore, EV-IR from miR34-knockout cells could not induce ROS and raised γH2AX to lesser extent than EV-IR from miR34-wild type cells. These results establish a novel role for the Abl-miR-34c DDR pathway in stimulating radiation-induced bystander effects.

## INTRODUCTION

In multicellular organisms, ionizing radiation (IR) causes breakage of cellular DNA to activate a wide range of responses not only in directly irradiated cells but also in neighboring or distant cells not targeted by IR (Mukherjee et al., 2014; Prise and O’Sullivan, 2009; Verma and Tiku, 2017). The non-target, or bystander, effects of IR occur when irradiated cells secrete soluble factors and/or extracellular vesicles (EV) to propagate the damage signal to naïve, non-irradiated cells (Jelonek et al., 2016; Mukherjee et al., 2014; Prise and O'Sullivan, 2009; Verma and Tiku, 2017). The master regulators of DNA damage response (DDR), i.e., ATM and p53, are required for irradiated cells to secrete bystander effectors (Burdak-Rothkamm et al., 2008; Komarova et al., 1998); however, how other DDR pathways stimulate the bystander effects of radiation is mostly unknown.

Previous studies have established that IR stimulates nuclear Abl tyrosine kinase to regulate transcription, DNA repair and microRNA processing (Baskaran et al., 1997; Kaidi and Jackson, 2013; Preyer et al., 2007; Shaul and Ben-Yehoyada, 2005; Tu et al., 2015; Wang, 2014). The ubiquitously expressed Abl has many context-dependent biological functions that are determined by its activating signals, its interacting proteins and its subcellular localization (Wang, 2014). Because DNA damage signal initiates in the nucleus, we investigate how nuclear Abl regulates DDR. Towards this goal, we mutated the three nuclear localization signals (NLS) in the mouse *Abl1* gene to create the *Abl-µNLS* (*µ*) allele (Preyer et al., 2007). We found that cisplatin-induced apoptosis was reduced in the *Abl^µ/µ^* embryonic stem cells and in the renal proximal tubule epithelial cells (RPTC) of the *Abl^µ/µ^* mice (Preyer et al., 2007; Sridevi et al., 2013), providing *in vivo* confirmation for the *in vitro* finding that cisplatin activates Abl to stimulate p73-mediated and p53-independent apoptosis in human colon cancer cells (Gong et al., 1999). To identify other nuclear Abl-stimulated pro-apoptotic factors, we searched for and found that Abl kinase stimulates the processing of precursor miR-34c, and that the induction of miR-34c by cisplatin is defective in *Abl-µNLS* mice (Tu et al., 2015). The transcription of primary miR-34a and miR-34b/c is stimulated by p53 (He et al., 2007a); however, p53-dependent apoptotic response to DNA damage is not affected by the knockout of all three members (a, b, c) of the miR-34-family (Concepcion et al., 2012). These mouse genetics results propelled us to consider alternative functions for the Abl-miR34c pathway in DDR.

It has been shown that extracellular vesicles (EV) can transfer microRNAs between cells (Tkach and Thery, 2016; Valadi et al., 2007). Recent results have suggested that EV and microRNA are involved in the communication between irradiated and bystander cells (Chaudhry, 2014; Jelonek et al., 2016). Therefore, we investigated the role of nuclear Abl and miR-34c in EV-mediated bystander effects of radiation.

## RESULTS

### Isolation and Characterization of Extracellular Vesicles (Fig. 1)

We isolated extracellular vesicles (EV) by differential ultracentrifugation of media conditioned by non-irradiated (C) or irradiated (IR, 10 Gy) mouse embryo fibroblasts (MEFs) and avoided EV from serum by switching cells into serum-free media with 1% BSA before irradiation (Fig. 1A). The total protein content of EV-C and EV-IR was found to be comparable among several independent preparations, showing that IR did not affect the total yield of protein in the EV pellets (Fig. 1C). Nanoparticle tracking analyses showed similar size distributions and particle concentrations among EV preparations from media conditioned by non-irradiated (EV-C) or irradiated (EV-IR) MEFs (Fig. 1B). As the particles ranged from 50 nm to 300 nm in diameter, these EV preparations were likely to contain a mixture of micro-vesicles derived from different intracellular compartments (Cocucci et al., 2009). When added to naïve, non-irradiated responder MEFs, fluorescent-labeled EV-C and EV-IR were internalized by 98-100% of cells at 24 hours (Fig. 1D) and to comparable intracellular levels (Fig. 1E). Thus, the differential response of non-irradiated MEFs to EV-C and EV-IR was unlikely to be due to differential uptake of these vesicles.

**Figure 1.**
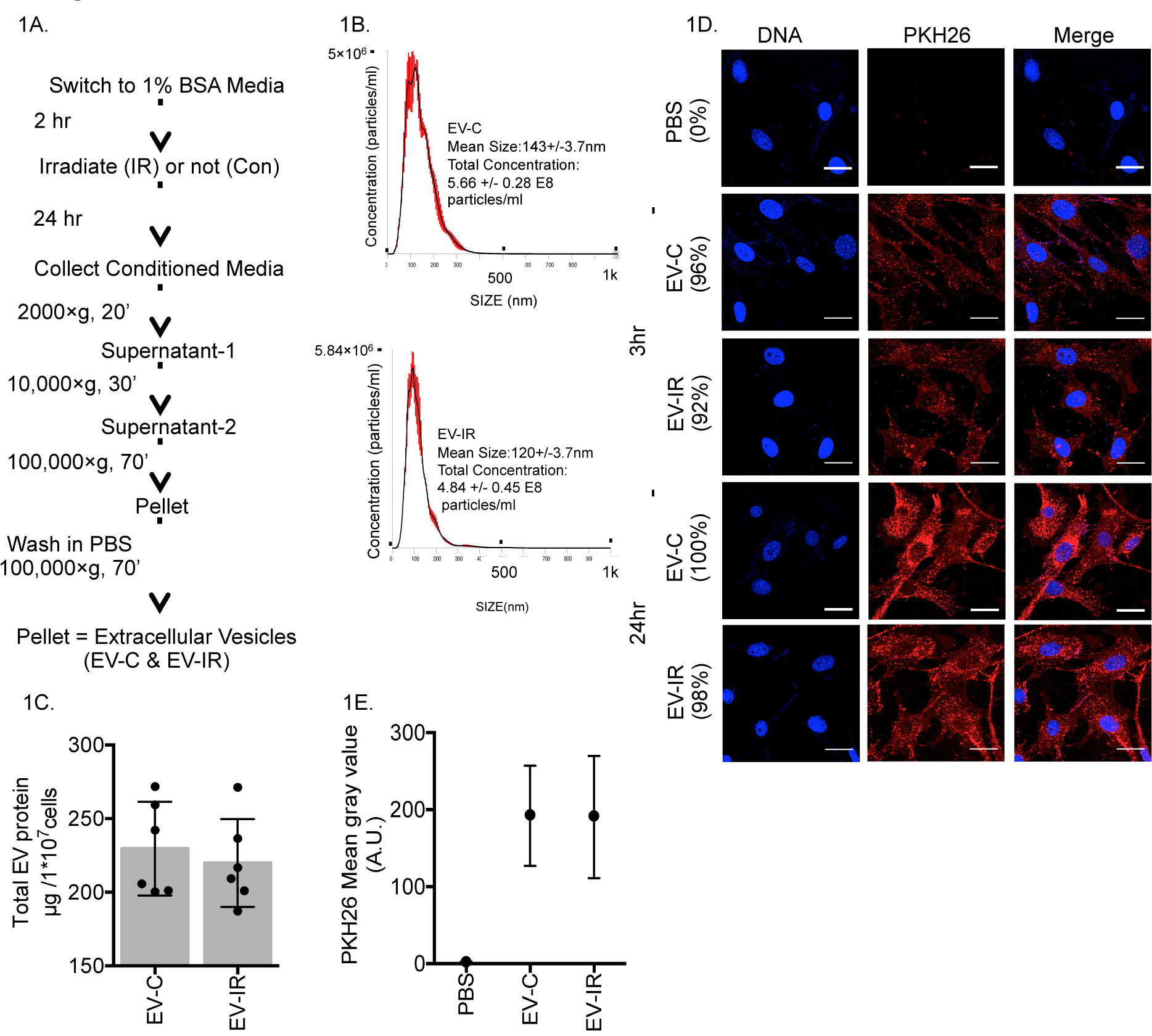
Isolation, Quantification and Uptake of Extracellular Vesicles. (A) Flow chart of EV isolation from media conditioned by non-irradiated (Con) or irradiated (IR, 10 Gy) mouse embryo fibroblasts. (B) Nanoparticle tracking analysis of a typical pair of EV-C & EV-IR preparations. (C) Total protein in EV-C or EV-IR preparations each from 10 million control or irradiated MEFs. Values shown are mean ± SD from six of each independent EV preparations. (D) Uptake of PKH26-labeled EV-C (25 μg) or EV-IR (25 μg) by naÏve, non-irradiated MEFs at 3 or 24 hours after EV addition. Representative fluorescence images of cells stained with Hoechst 33342 (DNA; blue) and EV stained with PKH26 dye (red) (Scale bar: 35 μm) with the percentage of PKH26-positive cells indicated. (E) Quantification of EV uptake in MEFs treated with PKH26-PBS, PKH26-EV-C (25 μg) or PKH26-EV-IR (25 μg) for 24 hours. The mean ± SD of PKH26 mean gray values from 200 cells are shown.

### EV-IR but not EV-C Inhibited Colony Formation (Fig. 2; Figs. S1, S2)

Inhibition of colony formation is both a direct and a bystander effect of ionizing radiation, for media conditioned by irradiated MEFs (CM-IR) inhibited colony formation when transferred to non-irradiated responder MEFs (Fig. 2A; Fig. S1A). We found that the EV-fraction retained the colony-inhibitory activity of CM-IR whereas the supernatant fraction lost most of that activity (Fig. 2A). Titration experiments showed that EV-IR inhibited colony formation in a dose-dependent manner, reaching saturation at a EV-protein level (25 µg) that was equivalent to several million particles per responder cell (Fig. 2C; Fig. S1C). By contrast, EV-C did not elicit such a dose-response (Fig. 2B; Fig. S1B). We then compared the growth inhibitory activity of IR vs. EV-IR. As expected, direct irradiation of MEFs induced p21Cip1 mRNA and protein (Fig. S2A-D), G2 arrest (Fig. S2E, F), and senescence (Fig. S2G, H). However, treatment with EV-IR did not induce p21Cip1 (Fig. S2A, B), G2 arrest (Fig. S2E, F), or senescence (Fig. S2G, H). These results showed that the colony-inhibitory mechanism of EV-IR differed from that of direct irradiation. Although immortal, the responder MEFs formed colonies at a low frequency of ~5%. It thus appeared that EV-IR inhibited a small fraction of colony-forming cells without causing growth arrest in the general population of responder cells that internalized EV-IR (Fig. 1D).

**Figure 2.**
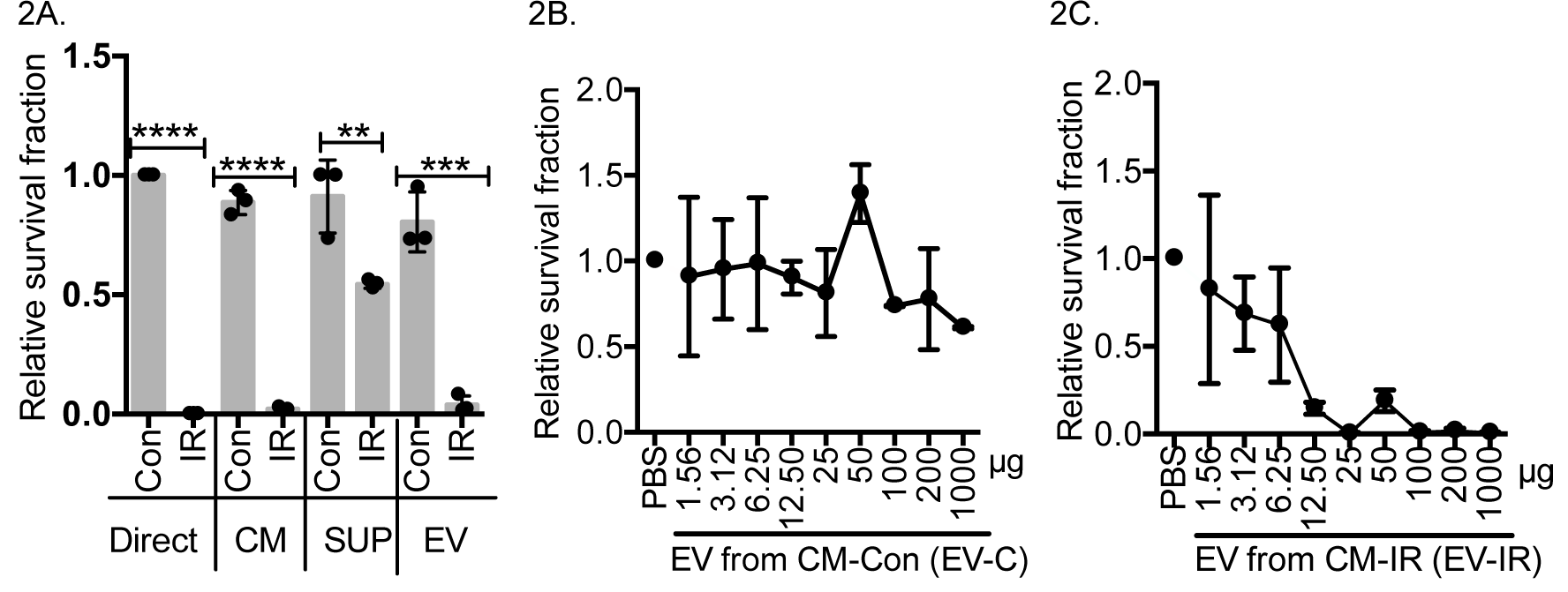
Extracellular Vesicles Secreted by Irradiated Cells Inhibited Colony Formation. (A) Survival fraction of MEFs at 15 days after the following treatments: non-irradiated (Direct, Con), 10 Gy irradiated (Direct, IR), treated for 24 hours with CM (conditioned media), Sup (supernatant fraction-2, Fig. 1A), or EV (washed extracellular vesicle pellet fraction, Fig. 1A, 25 μg each) from non-irradiated (Con) or 10 Gy irradiated (IR) MEFs. The survival fraction (number of colonies/number of cells seeded) of non-irradiated MEFs was set to 1. Values shown are mean ± SD from three independent experiments. \*\**P*<0.01, \*\*\**P*<0.001, \*\*\*\**P*< 0.0001, ONE WAY ANOVA. Representative images of colony formation results are shown in Fig S1A. (B) Survival fraction of MEFs at 15 days after treatment for 24 hours with PBS or the indicated concentrations of EV-C. The survival fraction of PBS-treated sample was set to 1. Values shown are mean ± SD from two independent experiments. Representative images of colony formation results are shown in Fig S1B. (C) Survival fraction of MEFs at 15 days after treatment for 24 hours with PBS or the indicated concentrations of EV-IR. The survival fraction of PBS-treated sample was set to 1. Values shown are mean ± SD from two independent experiments. Representative images of colony formation results are shown in Fig S1C.

### EV-IR but not EV-C Increased Reactive Oxygen Species (ROS) (Fig. 3; Fig. S3)

IR induces ROS in directly irradiated and non-irradiated bystander cells (Azzam et al., 2012; Klammer et al., 2015). We found that EV-IR, but not EV-C, dose-dependently increased the ROS levels in non-irradiated MEFs (Fig. 3A, B). This EV-IR-induced ROS occurred in the general population of responder cells (Fig. S3D) and was detectable at a EV-IR protein level (3.7 µg) that was equivalent to several hundred thousand particles per responder cell (Fig. 3B). The anti-oxidant N-acetyl-cysteine (NAC) neutralized EV-IR-induced ROS increase (Fig. 3A, EV-IR+NAC; Fig. S3D). NAC also reduced the colony inhibitory activity of EV-IR (Fig. 3C; Fig. S1D), suggesting that ROS was a contributing factor to EV-IR-induced inhibition of colony formation. Treatment with Proteinase K or RNase A did not abolish either the colony-inhibitory or the ROS-inducing activity of EV-IR (Fig. S3E), indicating that these effects were mediated by factors inside the vesicles.

**Figure 3.**
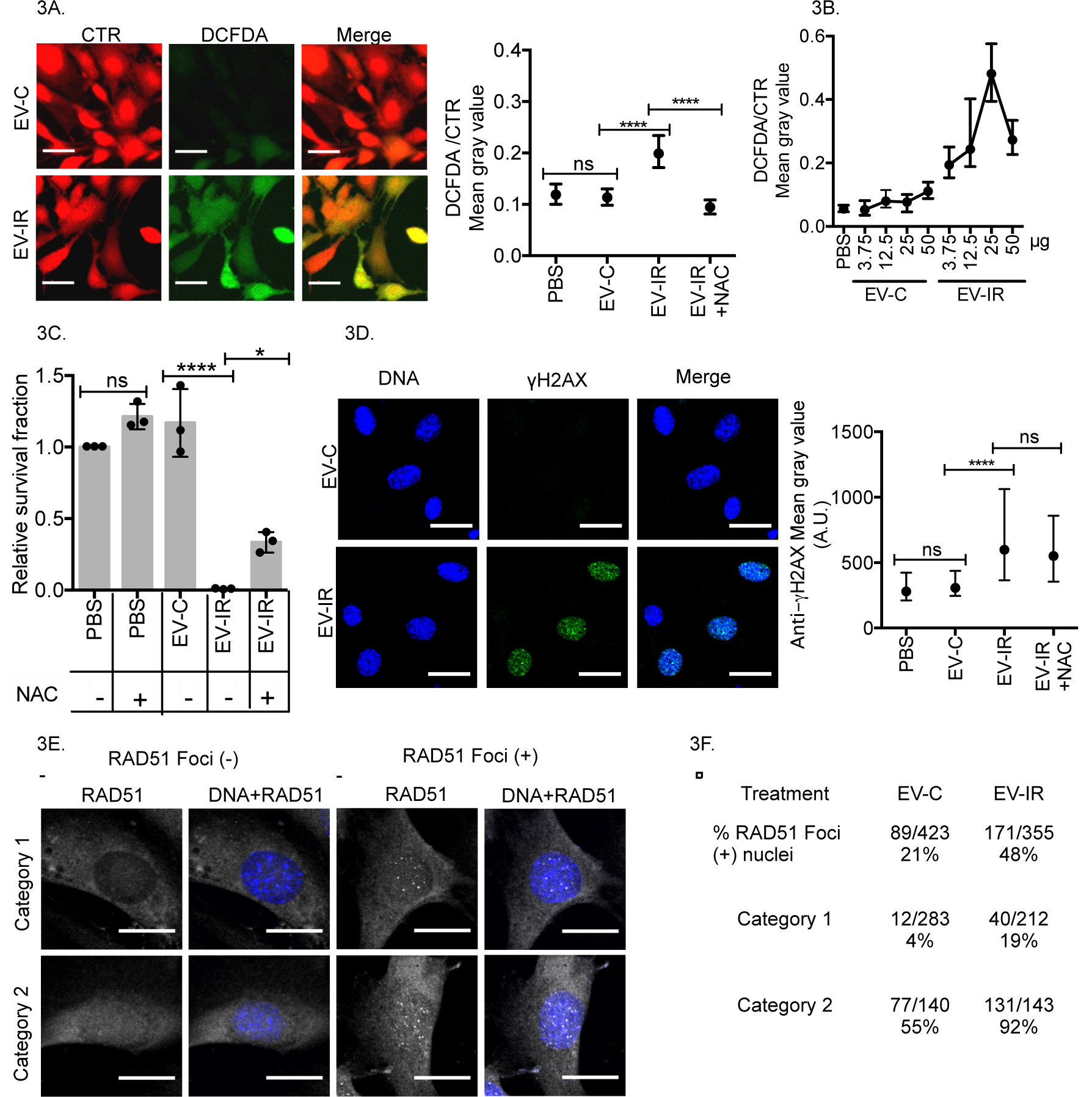
Extracellular Vesicles Secreted by Irradiated Cells Induced Reactive Oxygen Species & DNA Damage. (A) EV-IR but not EV-C increased ROS: live cells were stained with cell-tracker red (CTR) and DCFDA (green) at 24 hours after addition of EV-C, EV-IR or EV-IR+NAC (3.7 μg each of EV, 5 mM NAC) (Scale bar: 35μm). Ratio of DCFDA/CTR staining was determined as described in Methods. Values shown are medians with interquartile ranges from three independent experiments with at least 200 cells analyzed per sample per experiment. ns, not significant, \*\*\*\**P*≤ 0.0001, Kruskal-Wallis test. The DCFDA/CTR ratios of individual cells from one experiment are shown in Fig. S3D. (B) EV-IR dose-dependency in ROS induction: responder MEFs were treated with the indicated concentrations of EV-C or EV-IR for 24 hours and the ROS measured as in (A). Values shown are the medians and interquartile ranges of ratios from two independent experiments with at least 200 cells analyzed per sample per experiment. (C) NAC reduced the colony-inhibitory activity of EV-IR: Survival fraction of responder MEFs at 15 days after treatment for 24 hours with 25 μg each of EV-C, EV-IR, or EVIR+ NAC (5 mM). Relative survival fractions shown are mean ± SD from three independent experiments. ns, not significant, * *P*≤ 0.05, \*\*\*\**P*≤ 0.0001, ONE WAY ANOVA. Representative images of colonies are shown in Fig S1D. (D) EV-IR but not EV-C induced γH2AX foci: responder MEFs were fixed at 24 hours after addition of 3.7 μg each of EV-C, EV-IR or EV-IR+NAC (5 mM), stained with anti-γH2AX (green) & Hoechst 33342 (DNA; blue) (Scale bar: 35 μm) and the γH2AX levels quantified as described in Methods. Values shown are medians with interquartile ranges from three independent experiments with at least 200 cells analyzed per sample per experiment. ns, not significant, \*\*\*\**P*≤ 0.0001, Kruskal-Wallis test. The mean gray values of γH2AX in individual cells from one experiment are shown in Fig. S3C. (E) Scoring RAD51 foci: Responder cells were fixed and stained with anti-RAD51 (gray) & Hoechst 33342 (DNA; blue) at 24 hours after addition of EV-C or EV-IR (25 μg each) (Scale bar: 25 μm). Category-1 cells showed lower nuclear RAD51 signal than category-2 cells. (F) EV-IR but not EV-C increased RAD51 foci: Summary of RAD51 foci-positive nuclei in each indicated samples with breakdowns into categories. Representative images of multiple cells in each sample are shown in Fig. S3F. Summary of total nuclei scored per category is shown in Fig. S3G.

### EV-IR but not EV-C Increased γH2AX and RAD51 Foci (Fig. 3; Fig. S3)

Another bystander effect of radiation is the induction of DNA damage in non-irradiated cells (Klammer et al., 2010; Lorimore et al., 2003). We found that addition of EV-IR, but not EV-C, induced γH2AX foci in the non-irradiated responder MEFs (Fig. 3D; Fig. S3C). Quantification of images showed a range of γH2AX levels in MEFs (Fig. S3B, C). Treatment with EV-IR, but not EV-C, raised this range of γH2AX levels in the majority of responder cells (Fig. 3D; Fig. S3B, C). In side-by-side comparisons, we found that IR-induced γH2AX levels to be 2-fold higher than that induced by EV-IR (Fig. S3A, B). Although NAC blocked EV-IR-induced increase in ROS (Fig. 3A; Fig S3D), it did not block the EV-IR effect on γH2AX (Fig. 3D; Fig. S3C), indicating that ROS was not required for EV-IR to stimulate H2AX phosphorylation. Treatment with EV-IR also increased RAD51-foci in non-irradiated MEFs (Fig. 3E, F; Fig. S3F, G). We scored the RAD51-foci by visual inspection and separated the responder MEFs into two categories. The majority of responders (60-70%) showed lower nuclear RAD51-signal (category-1) (Fig. 3E; Fig. S3G); and among category-1 cells, 20% or 4% were positive for RAD51 foci after treatment with EV-IR or EV-C, respectively (Fig. 3F; Fig. S3F). Among category-2 cells that showed higher nuclear RAD51-signal (Fig. 3E; Fig. S3G), 90% or 55% were positive for RAD51-foci after treatment with EV-IR or EV-C, respectively (Fig. 3F; Fig. S3F). These results showed that EV-IR could induce bystander DNA damage in non-irradiated cells. Because EV-IR-induced increases in ROS and γH2AX occurred in a substantial population of responder cells that internalized these vesicles, we focused subsequent studies on these two bystander effects.

### Nuclear Abl not Required for Radiation to Induce ROS and γH2AX (Fig. 4)

To determine the essential function of nuclear Abl in DDR, we constructed the *Abl-µNLS* allele in the mouse *Abl1* gene by mutating the three nuclear localization signals (NLS) in the Abl protein (Fig. 4A) (Preyer et al., 2007). We then established embryo fibroblasts (MEFs) from littermate *Abl^+/+^* (*Abl-wt*) and *Abl^µ/µ^* (*Abl-µNLS*) mice through serial passages in culture. Irradiation of *Abl-wt* MEFs induced nuclear accumulation of Abl, whereas irradiation of *Abl-µNLS* MEFs did not induce nuclear accumulation of Abl-µNLS (Fig. 4B). Thus, mutation of the NLS is sufficient to abolish IR-induced Abl nuclear accumulation. However, IR still increased ROS and γH2AX in the *Abl-µNLS* MEFs (Fig. 4C-F), showing that nuclear Abl is not required for IR to cause these effects in directly irradiated MEFs.

**Figure 4.**
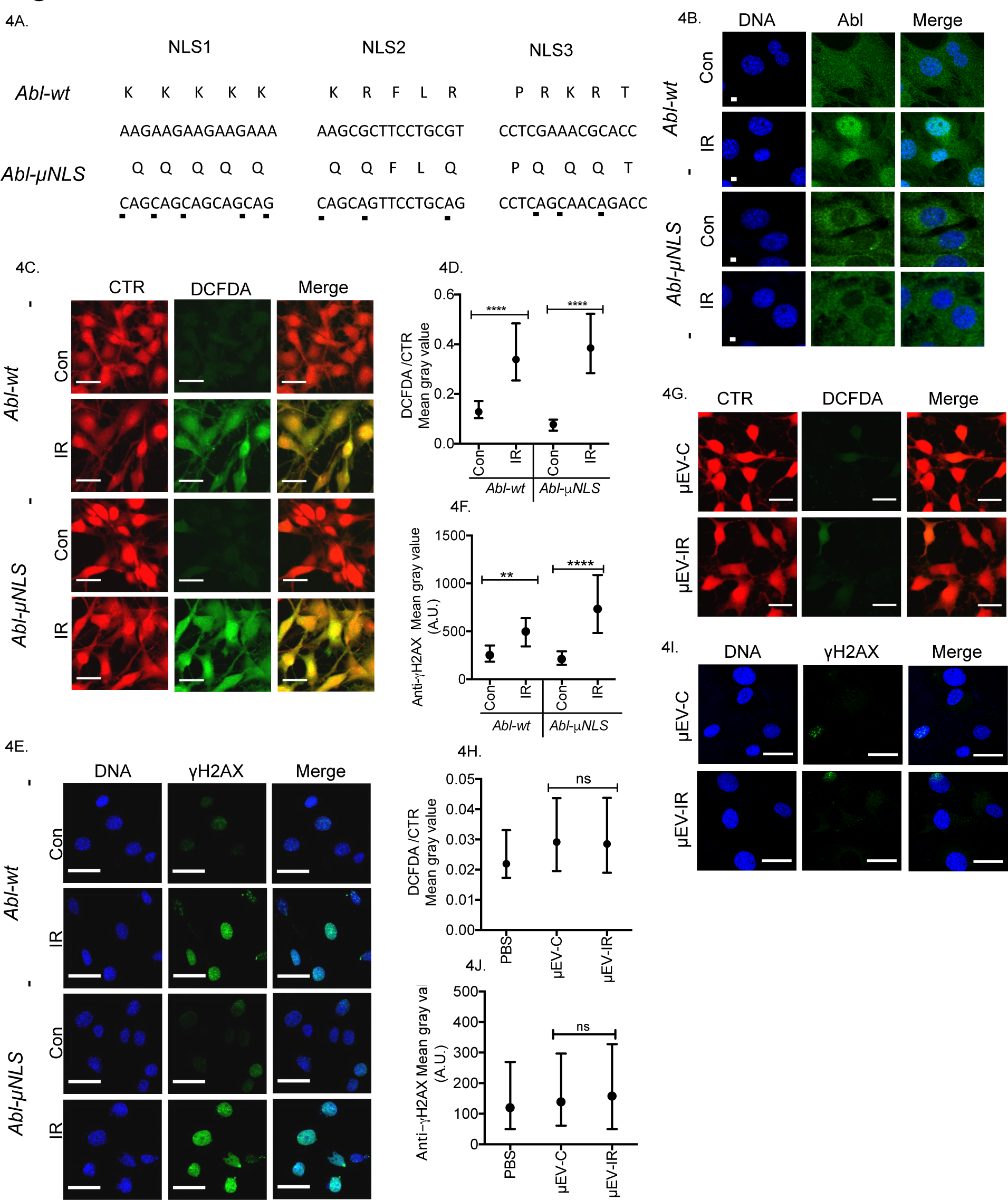
Extracellular Vesicles Secreted by Irradiated *Abl-μNLS* Cells Failed to Induce ROS & γH2AX. (A) Substitution mutations of the three Abl nuclear localization signals (NLS) in the *Abl-μNLS* allele. (B) Representative images of anti-Abl (green) and Hoechst 33342 (DNA; blue) staining in the indicated MEFs: Con, no irradiation; IR: 3 hours after 10 Gy (Scale bar: 30 μm). (C & D) Direct irradiation induced ROS in *Abl-μNLS* MEFs: the indicated cells were stained with CTR (red) and DCFDA (green) at 24 hours after no irradiation (Con) or 10 Gy of IR (Scale bar: 35 μm). Values shown are the medians with interquartile ranges of DCFDA/CTR ratios from two independent experiments with at least 200 cells analyzed per sample per experiment. **** *P*≤ 0.0001, Kruskal-Wallis test. (E & F) Direct irradiation increased γH2AX in Abl-γNLS MEFs: the indicated cells were fixed and stained with anti-γH2AX (green) and Hoechst 33342 (DNA; blue) at 24 hours after no irradiation or 10 Gy IR (Scale bar: 35 μm). Values shown are medians with interquartile ranges of anti-γH2AX mean gray values from two independent experiments with at least 200 cells analyzed per sample per experiment. ** *P*≤ 0.01, **** *P*≤ 0.0001, Kruskal-Wallis test. (G & H) γEV-IR did not increase ROS: responder MEFs (*Abl-wt*) stained with CTR (red) and DCFDA (green) at 24 hours after treatment with γEV-C & μEV-IR (3.7 μg each) isolated from CM of non-irradiated or irradiated (10 Gy IR) Abl-γNLS MEFs (Scale bar: 35 μm). Values shown are medians with interquartile ranges of DCFDA/CTR ratios from three independent experiments with at least 200 cells analyzed per sample per experiment. ns: not significant, Kruskal-Wallis test (I & J) γEV-IR did not increase γH2AX: responder MEFs (*Abl-wt*) were fixed and stained at 24 hours after treatment with γEV-C & γEV-IR (3.7 μg each) (Scale bar: 35 μm). Values shown are medians with interquartile ranges of anti-γH2AX mean gray values from three independent experiments with at least 200 cells analyzed per sample per experiment. ns, not significant, Kruskal-Wallis test.

### Nuclear Abl Required for ROS- and γH2AX-Inducing Activities of EV-IR (Figs. 4, 5, 6)

Although IR induced ROS and γH2AX in directly irradiated *Abl-µNLS* MEFs, we found that EV from irradiated *Abl-µNLS* MEFs (µEV-IR) did not induce ROS (Fig. 4G, H) or increase γH2AX (Fig. 4I, J) in responder MEFs (*Abl-wt)*. To determine if restoration of nuclear Abl could rescue these µEV-IR defects, we stably expressed Abl^WT^ or Abl^µNLS^ proteins in *Abl-µNLS* MEFs through retroviral-mediated gene transfer (Fig. 5A) without significantly raising the overall levels of Abl protein (Fig. 5B). After irradiation (10 Gy IR), we found nuclear accumulation of Abl^WT^ but not Abl^µNLS^ in the *Abl-µNLS* MEFs (Fig. 5C). The expression of Abl^WT^ or Abl^µNLS^ did not alter the direct effects of IR in the *Abl-µNLS* MEFs (Fig. 5D-G), again showing that nuclear Abl did not make significant contributions to ROS and γH2AX in irradiated MEFs. However, expression of Abl^WT^ but not Abl^µNLS^ restored the ROS-and γH2AX-inducing activities of µEV-IR (Fig. 6). Together, these results establish that nuclear entry of Abl in irradiated cells is required for the formation of EV-IR with ROS-and γH2AX-inducing activities.

**Figure 5.**
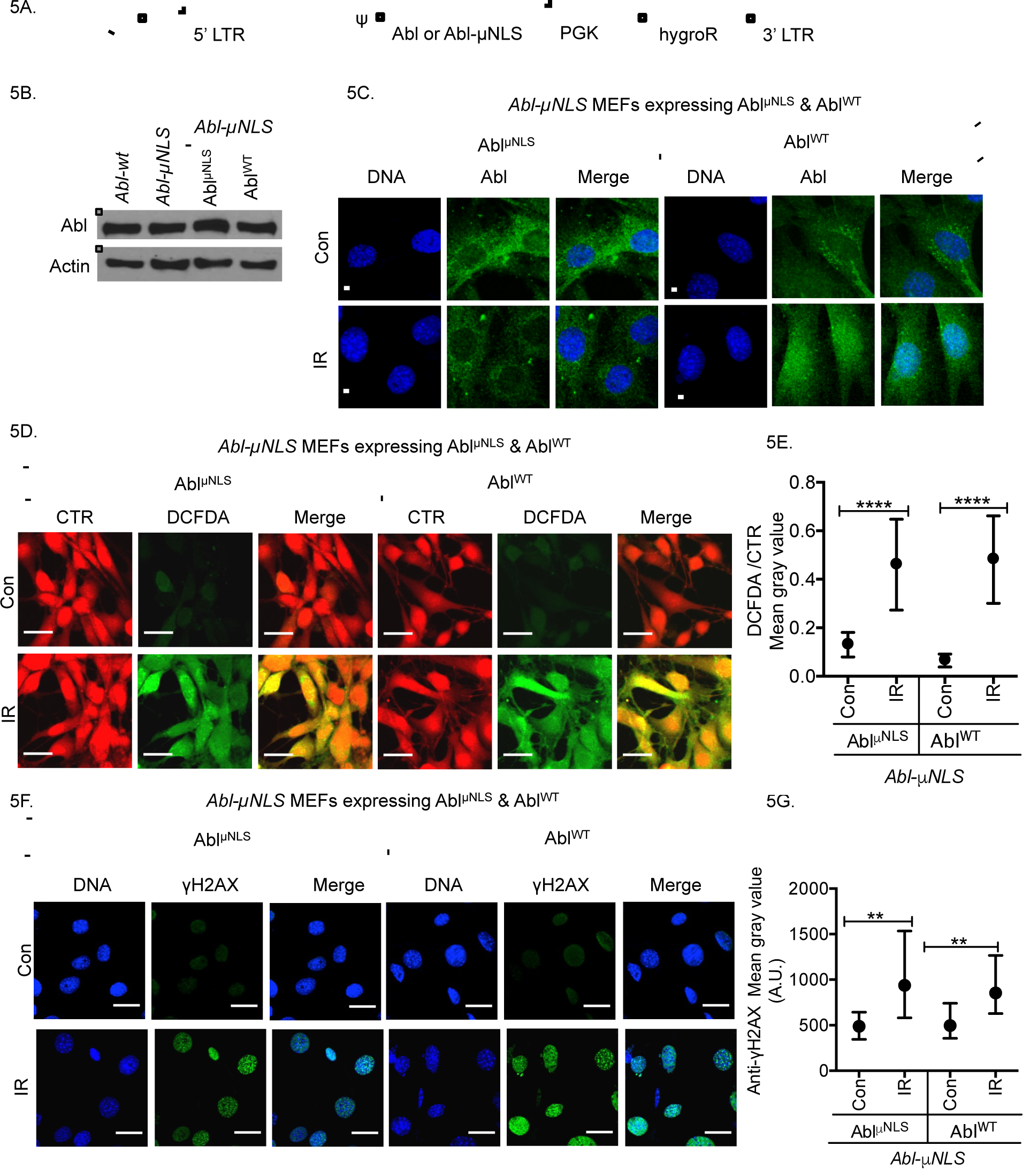
Radiation-induced ROS and γH2AX in *Abl-μNLS* Cells Expressing Abl^WT^ or Abl^μNLS^. (A) pMSCV retroviral constructs for expression of Abl^WT^ or Abl^γNLS^. (B) Immunoblotting of Abl in whole lysates of the indicated cells. (C) Representative fluorescence images at 3 hours after irradiation (10 Gy) in the indicated MEFs. Hoechst 33342 (DNA; blue), anti-Abl (green) (Scale bar: 30 μm). (D) Representative images & quantification of ROS: the indicated live cells were stained with CTR (red), and DCFDA (green) at 24 hours post irradiation (10 Gy) (Scale bar: 35 μm). Values shown are the medians with interquartile ranges of DCFDA/CTR rations from two independent experiments with at least 200 cells analyzed per sample per experiment. **** *P*≤ 0.0001, Kruskal-Wallis test. (E) Representative images & quantification of γH2AX: Hoechst 33342 (DNA; blue), anti-γH2AX (green), (Scale bar: 35 μm). Values shown are medians with interquartile ranges of anti-γH2AX mean gray values from two independent experiments with at least 200 cells analyzed per sample per experiment. ** *P*≤ 0.01, Kruskal-Wallis test.

**Figure 6.**
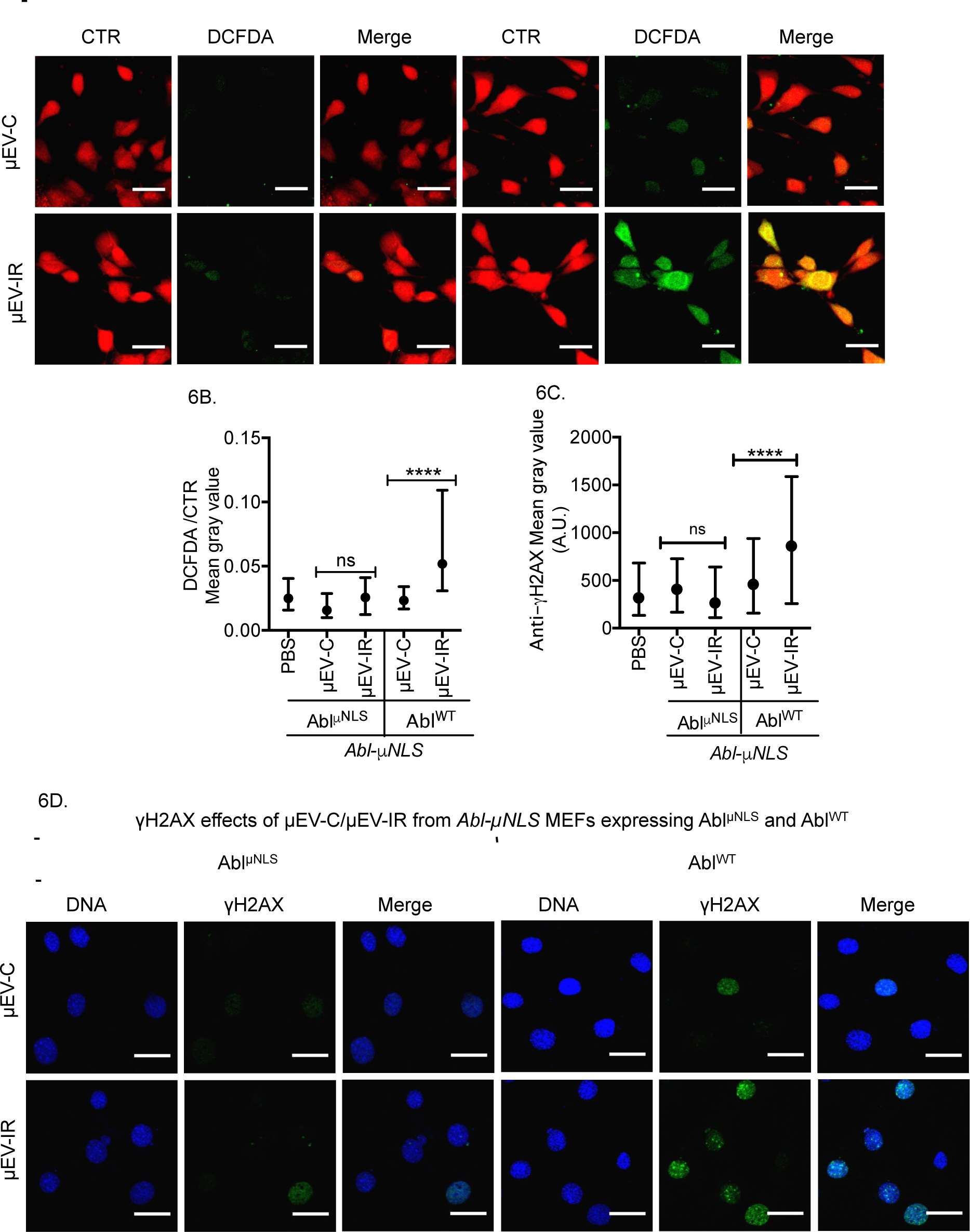
Expression of Abl^WT^ but not Abl^μNLS^ Restored ROS- and γH2AX-Inducing Activities of μEV-IR. (A & B) Representative images & quantification of ROS: live responder cells were stained with CTR (red) and DCFDA (green) at 24 hours after treatments with the indicated EV (3.7 μg each) (Scale bar: 35 μm). Values shown are medians with interquartile ranges of DCFDA/CTR ratio from three independent experiments with at least 200 cells analyzed per sample per experiment. ns: not significant, \*\*\*\**P*≤ 0.0001, Kruskal Wallis test. (C & D) Representative images & quantification of γH2AX: responder cells treated with the indicated EV (3.7 μg each) were fixed and stained at 24 hours after EV addition. Hoechst 33324 (DNA; blue), anti-γH2AX (green) (Scale bar: 35 μm). Values shown are medians with interquartile ranges of anti-γH2AX mean gray values from three independent experiments with at least 200 cells analyzed per sample per experiment. ns: not significant, \*\*\*\**P*≤ 0.0001, Kruskal-Wallis test.

### Abl Kinase Inhibitor Abolished the ROS- and γH2AX-Inducing Activities of EV-IR (Fig. 7)

Ionizing radiation not only induces Abl nuclear accumulation but it also activates Abl kinase activity (Baskaran et al., 1997; Kaidi and Jackson, 2013). To assess the role of Abl kinase, we pre-treated MEFs with the Abl kinase inhibitor imatinib before irradiation (Fig. 7A) and compared the bystander effects of EV-IR with EV-(IM+IR). We found that the ROS-and γH2AX-inducing activities of EV-(IM+IR) were significantly reduced when compared to EV-IR (Fig. 7B-E). Thus, Abl kinase activity is also required for irradiated cells to produce EV-IR with ROS-and γH2AX-inducing activities.

**Figure 7.**
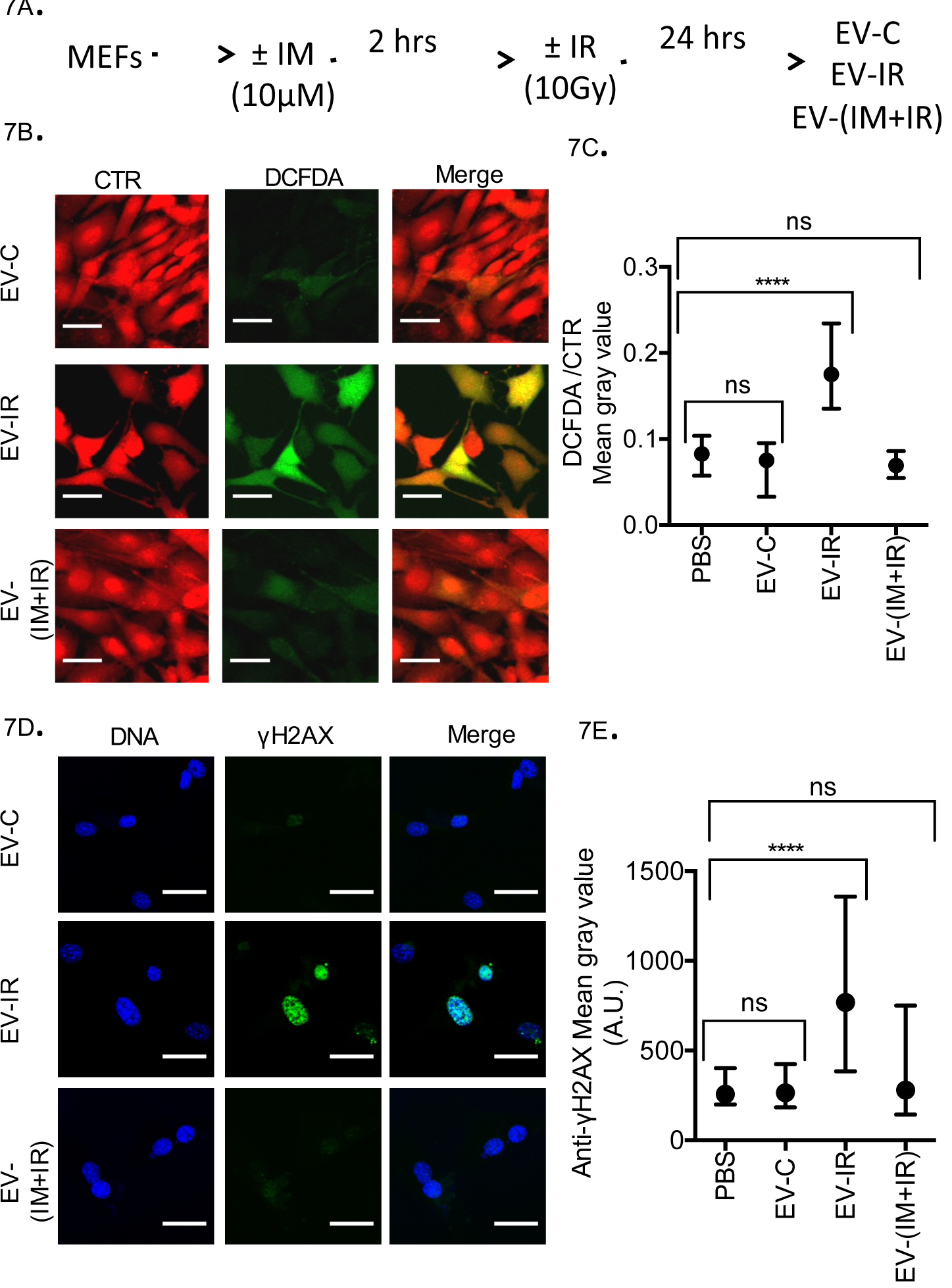
Treatment of Irradiated Cells with Abl Kinase Inhibitor Abolished the ROS- and γH2AX-Inducing Activities of Extracellular Vesicles. (A) Timeline of experimental protocol. (B & C) Representative images & quantification of ROS: live responder cells treated with the indicated EV were stained with CTR (red) and DCFDA (green) at 24 hours after EV addition (Scale bar: 35 μm). Values shown are medians with interquartile ranges of DCFDA/CTR ratios from three independent experiments with at least 200 cells analyzed per sample per experiment. ns, not significant, **** *P*≤ 0.0001, Kruskal-Wallis test. (D & E) Representative images & quantification for γH2AX: responder cells were fixed at 24 hours after treatments with the indicated EV and stained with Hoechst 33324 (DNA; blue) and anti-γH2AX (green) (Scale bar: 30 μm). Values shown are medians with interquartile ranges of anti-γH2AX mean gray values from three independent experiments with at least 200 cells analyzed per sample experiment. \*\*\**P*≤ 0.001, **** *P*≤ 0.0001, Kruskal-Wallis test.

### Nuclear Abl Raised miR-34c Levels in EV-IR for Transfer into Responder Cells (Fig. 8)

IR stimulates the expression of many miRs in directly irradiated cells (Chaudhry, 2014; He et al., 2007a). We have previously shown that Abl kinase stimulates the processing of *pri-miR34b/c* to *pre-miR-34b* and *pre-miR-34c* (Tu et al., 2015). Since the majority of intracellular miRs are found in EV (Shurtleff et al., 2016), we measured the levels of miR-34c in MEFs and in EV. With *Abl-wt* MEFs, irradiation increased the intracellular miR-34c by 3-fold (Fig. 8A), and the EV-IR miR-34c by 20-fold (Fig. 8D). With *Abl-µNLS* MEFs, radiation increased the intracellular miR-34c by 2-fold (Fig. 8B). However, this 2-fold intracellular increase did not raise the miR-34c levels in µEV-IR (Fig. 8E). In the responder MEFs, we found a 2-fold increase in miR-34c after treatment with EV-IR, but not with EV-C, µEV-C or µEV-IR (Fig. 8G, H). Expression of Abl^WT^ in *Abl-µNLS* MEFs raised IR-induced miR-34c levels (Fig. 8C) and restored the miR-34c increase in µEV-IR by 2-fold (Fig. 8F). Although the rescue of miR-34c increase in µEV-IR by Abl^WT^ did not reach the 20-fold level found with EV-IR, treatment with µEV-IR from Abl^WT^ expressing *Abl-µNLS* cells did cause a 2-fold increase in the intracellular miR-34c levels in responder MEFs (Fig. 8I). We also measured the mRNA levels of three previously confirmed miR-34c target genes (Cannell et al., 2010; Garofalo et al., 2013), namely *Pdgfra, Pdgfrb* and *Myc*, in responder cells after treatment with EV-C or EV-IR. We found that EV-IR treatment reduced *Myc* RNA in responder cells, indicating that the miR-34c transferred by EV-IR was functional in targeting *Myc* but not *Pdgfra* or *Pdgfrb* for down-regulation (Fig. 8J). Together, these results show that nuclear Abl not only contributes to IR-induced miR-34c expression, but also drives the secretion of functional miR-34c in EV-IR for transfer into responder cells.

**Figure 8.**
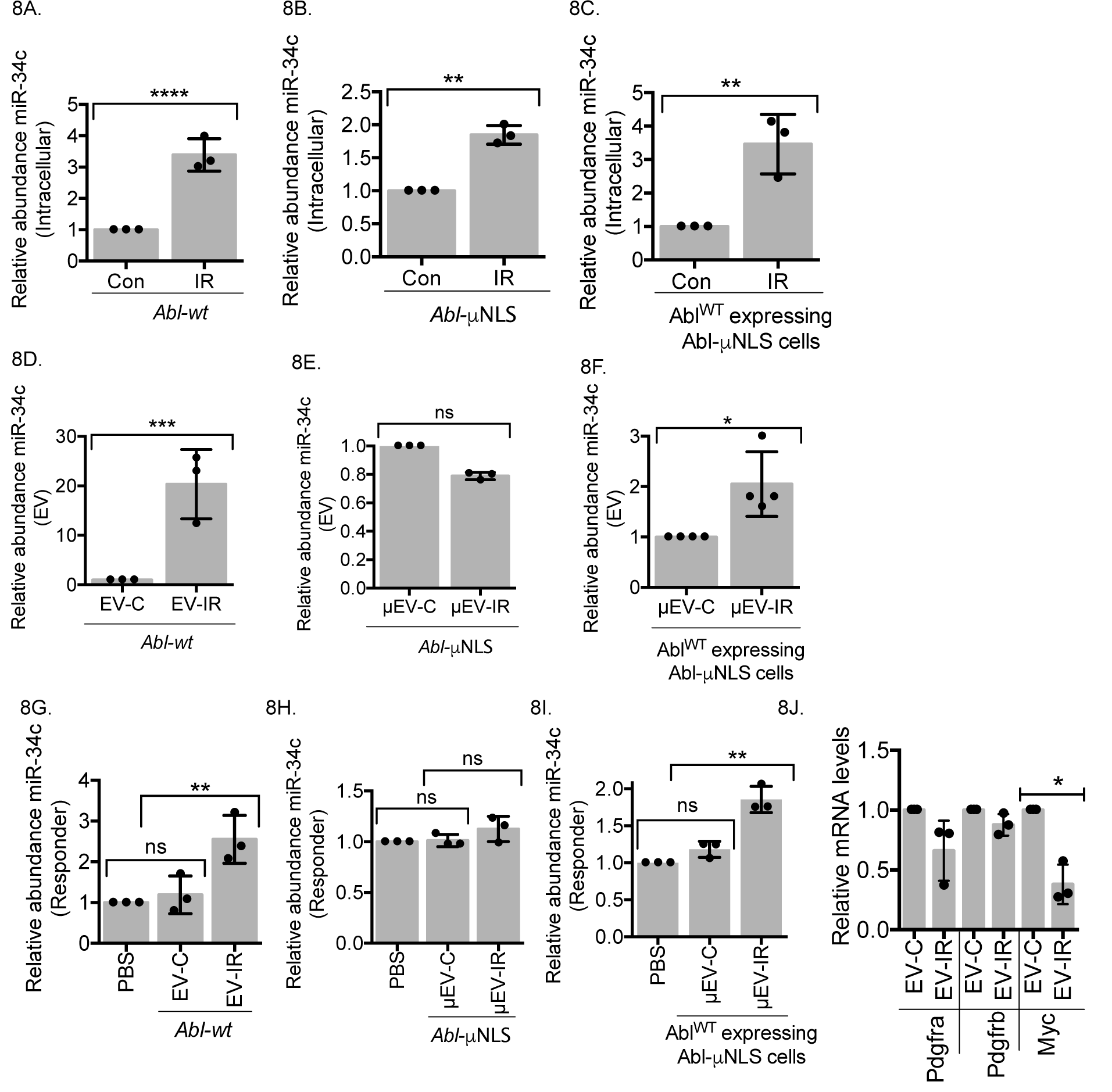
Nuclear Abl Required for Raising miR-34c Levels in Extracellular Vesicles from Irradiated Cells for Transfer into Responder Cells. (A, B, C) Irradiation increased intracellular levels of miR-34c: *Abl-wt* (A), *Abl-μNLS* (B) and Abl^WT^-expressing *Abl-μNLS* MEFs (C) were irradiated at 10 Gy, RNA was collected after 24 hours, and miR-34c measured as described in Methods. Abundance of miR-34c normalized to that of U6 in each non-irradiated (Con) MEFs was set to 1. Values shown are relative miR-34c abundance (mean ± SD) from three independent experiments. ** *P*≤ 0.01, \*\*\*\**P*≤ 0.0001, ONE WAY ANOVA. (D, E, F) Irradiation-induced miR-34c increase in EV required nuclear Abl: Relative abundance of miR-34c in EV isolated from CM of non-irradiated or irradiated (IR, 10 Gy) *Abl-wt* (D), *Abl-μNLS* (E) or Abl^WT^-expressing *Abl-μNLS* MEFs (F). Values shown are relative abundance (mean ± SD) from three independent experiments with U6- normalized miR-34c in EV-C, μEV-C, or μEV-C(Abl^WT^) set to 1. ns, not significant, *P. 0.05,\*\*\**P*≤ 0.001, ONE WAY ANOVA. (G, H, I) Treatment with miR-34c-containing EV raised the levels in responder cells: responder cells (non-irradiated *Abl-wt* MEFs) treated with PBS or the indicated EV (25 μg) from (D, E, F) were harvested at 24 hours and intracellular miR-34c measured as described in Methods. The U6-normalized miR-34c abundance in PBS-treated cells was set to 1. Values shown are relative abundance (mean ± SD) from three independent experiments. ns: not significant, \*\**P*≤ 0.01, ONE WAY ANOVA. (J) Treatment with EV-IR but not EV-C reduced Myc RNA in responder cells: Relative RNA levels of the indicated genes after treatment with the indicated EV (25 μg each) for 24 hours. Values shown are relative RNA abundance (mean ± SD) from three independent experiments. \**P*≤ 0.05, ONE WAY ANOVA.

### Ectopically Produced EV-miR34c with ROS-and γH2AX-Inducing Activities (Fig. 9; Fig. S4)

To ectopically produce EV with miR-34c from non-irradiated cells, we transfected HEK293T cells with a miR-34c-minigene and a constitutively activated Abl kinase (AblPPn) (Fig. 9A, B). The miR-34c-minigene raised miR-34c levels in transfected cells (Fig. 9C) and in EV isolated from the media of those transfected cells (Fig. 9D). Co-expression with AblPPn further increased the intracellular and the EV levels of miR-34c (Fig. 9C, D). When added to responder MEFs, EV-miR-34c and EV-miR-34c+AblPPn increased the intracellular levels of miR-34c proportional to miR-34c levels in the EVs (Fig. 9E). Furthermore, EV-miR-34c and EV-miR-34c+AblPPn increased the ROS and the γH2AX levels in responder MEFs proportional to miR-34c levels (Fig. 9F, G, Fig. S4A, B, C, D). These results showed that ectopically expressed miR-34c is secreted in EV by HEK293T cells, that activated Abl kinase stimulates the EV-levels of miR-34c, that EV-miR34c transfers miR-34c into responder cells, and that the levels of miR-34c correlate with the levels of ROS and γH2AX in responder cells.

**Figure 9.**
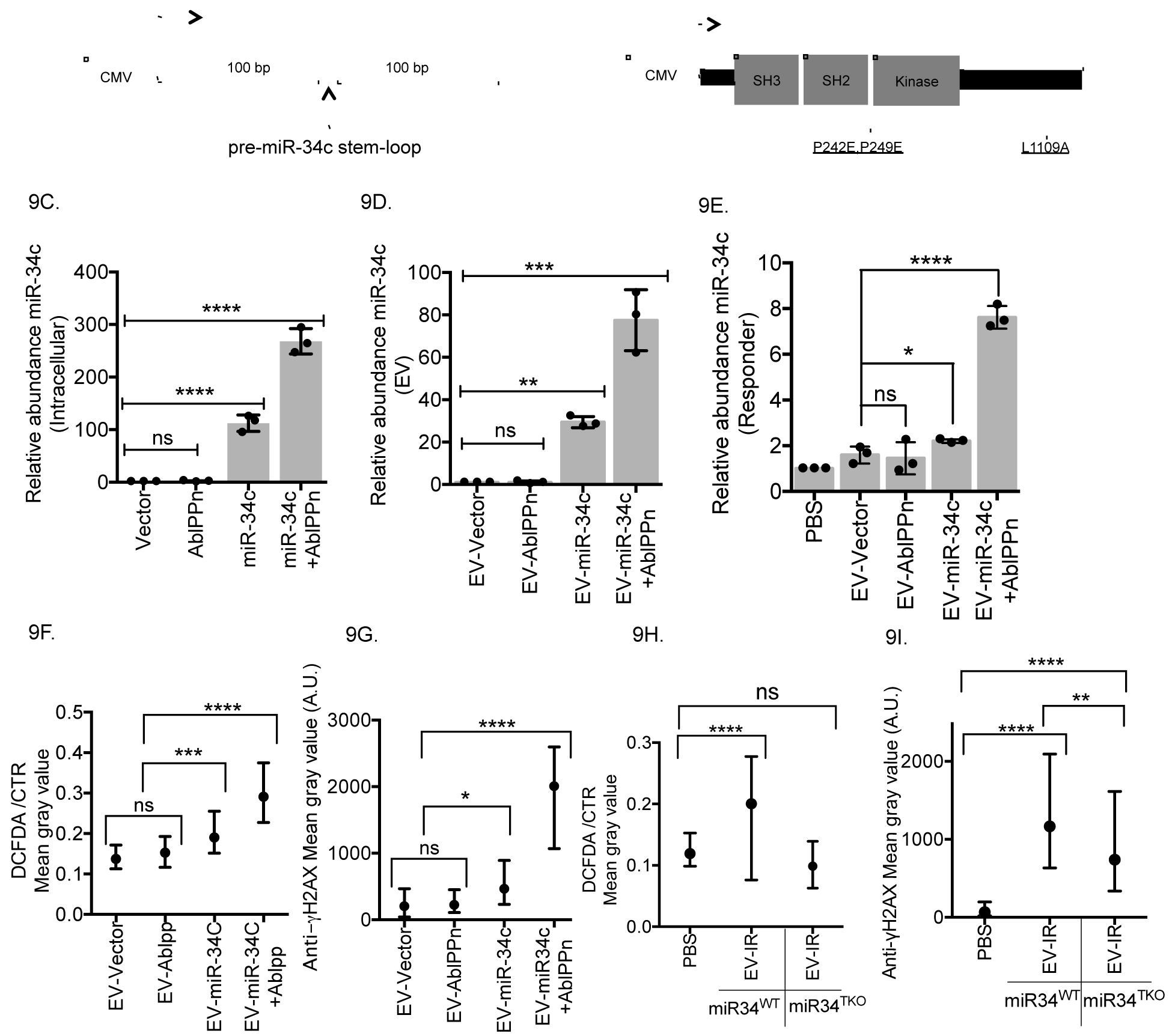
Induction of ROS and γH2AX by Extracellular Vesicles with miR-34c. (A, B) The miR-34c minigene and the AblPPn expression plasmids. (C, D) Co-transfection of miR-34c-minigene with AblPPn raised intracellular (C) and EV (D) miR-34c levels: HEK293T cells transfected with vector, AblPPn, miR-34c-minigene or miR-34c-minigene+AblPPn were collected at 24 hours post transfection and the relative abundance of miR-34c measured in cells (C) and in EV isolated from media of the transfected cells (D). U6-normalized miR-34c abundance in vector-transfected cells (C) or vector-transfected EV (D) was set to 1. Values shown are mean ± SD from three independent experiments. ns: not significant, \*\**P*≤ 0.01, \*\*\**P*≤ 0.001, \*\*\*\**P*≤ 0.0001, ONE WAY ANOVA. (E) Treatment with miR-34c-containing EV raised miR-34c levels in responder MEFs: non-irradiated MEFs (*Abl-wt*) were treated with PBS or the indicated EV preparations (25 μg each) for 24 hours. U6-normalized miR-34c abundance in PBS-treated MEFs was set to 1. Values shown are mean ± SD from three independent experiments. ns: not significant, \**P*≤ 0.05, \*\*\*\**P*≤ 0.0001, ONE WAY ANOVA. (F) miR-34c-containing EV induced ROS: DCFDA/CTR ratios in responder MEFs at 24 hours after treatment with the indicated EV (25 μg each) from the transfected HEK239T cells. Values shown are medians with interquartile ranges from three independent experiments with at least 200 cells analyzed per sample per experiment. ns: not significant, \*\*\**P*≤ 0.001, \*\*\*\**P*≤ 0.0001, Kruskal Wallis test. Representative images and the ratios of individual cells from one experiment are shown in Fig. S4A, C. (G) miR-34c-containing EV increased γH2AX: Levels of γH2AX in responder MEFs at 24 hours after treatment with the indicated EV (25 μg each) from the transfected HEK239T cells. Values shown are medians with interquartile ranges from two independent experiments with at least 200 cells analyzed per sample per experiment. ns: not significant, \**P*≤ 0.05,\*\*\*\**P*≤ 0.0001, Kruskal-Wallis test. Representative images and the mean gray values of individual cells from one experiment are shown in Fig. S4B, D. (H) EV-IR from miR34TKO MEFs failed to induce ROS: DCFDA/CTR ratios in responder MEFs (*Abl-wt*) treated with PBS or the indicated EV-IR (25 μg each) isolated from irradiated miR34WT or miR34TKO MEFs. Values shown are medians with interquartile ranges from two independent experiments with at least 200 cells analyzed per sample per experiment. ns: not significant, ****P. 0.0001, Kruskal-Wallis test. Representative images and the ratios of individual cells from one experiment are shown in Fig. S4 E, G. (I) EV-IR from miR34^TKO^ cells showed reduced γH2AX induction: Levels of γH2AX in responder MEFs (*Abl-wt*) treated with PBS or the indicated EV-IR (25 μg each) isolated from irradiated miR34^WT^ or miR34^TKO^ MEFs. Values shown are medians with interquartile ranges from two independent experiments with at least 200 cells analyzed per sample per experiment. ** *P*≤ 0.01 \*\*\*\**P*≤ 0.0001, Kruskal-Wallis test. Representative images and the mean gray values of individual cells in each experimental sample are shown in supplementary Fig. S4F, H.

### Defects of EV-IR from miR-34-Triple Knockout MEFs in Inducing ROS and γH2AX (Fig. 9; Fig. S4)

To determine if miR-34c is necessary for EV-IR to induce ROS and γH2AX, we isolated EV-IR-miR34^TKO^ from media conditioned by irradiated primary MEFs derived from the miR34-family (a, b, c) triple knockout mice, and EV-IR-miR34^WT^ from irradiated primary MEFs derived from littermate wild-type mice (Concepcion et al., 2012). We found that EV-IR-miR34^WT^ induced ROS in the responder MEFs, showing that EV-IR from primary (miR34^WT^) and established (*Abl-wt*) MEFs had similar ROS-inducing activity. However, EV-IR-miR34^TKO^ did not induce ROS in responder MEFs (Fig. 9H; Fig. S4E, G). By contrast, EV-IR-miR34^TKO^ was able to cause γH2AX increase in responder cells, but to a significantly lower level than that caused by EV-IR-miR34^WT^(Fig. 9I; Fig.S4 F,H). These results suggest that miR-34-family is required for the ROS-inducing activity of EV-IR; and that this family of microRNAs contribute to, but are not the only inducers of, γH2AX increase in non-irradiated bystanders.

### Roscovitine Inhibited EV-IR-Induced γH2AX (Fig. 10; Fig. S5)

The induction of bystander DNA damage is a detrimental side effect of radiation therapy as it is associated with secondary malignancy (Burtt et al., 2016; Lorimore et al., 2008). Previous studies have linked the induction of bystander DNA damage to oxidative stress (Havaki et al., 2015). However, we found that NAC neutralized EV-IR-induced ROS without affecting the increase in γH2AX (Fig. 3). Because EV-IR stimulated γH2AX and RAD51 foci in only a subpopulation of responder cells (Fig. 3; Fig. S3), and because γH2AX and RAD51 foci can occur during DNA replication (Scully et al., 1997; Tashiro et al., 1996), we inhibited DNA replication by treating synchronized populations of responder cells with the Cdk-inhibitor Roscovitine (Rosc) (Fig. 10A). In the absence of Rosc, treatment of synchronized responders with EV-IR again caused an increase in γH2AX (Fig. 10B; Fig. S5A, G). However, in the presence of Rosc, EV-IR failed to increase γH2AX (Fig. 10B; Fig. S5A, G). Similar results were obtained with EV-miR-34c isolated from miR-34c-minigene transfected HEK293T cells (Fig. 10C; Fig. S5B, H). Thus, EV-IR and EV-miR-34c might cause replication stress to induce bystander DNA damage.

**Figure 10.**
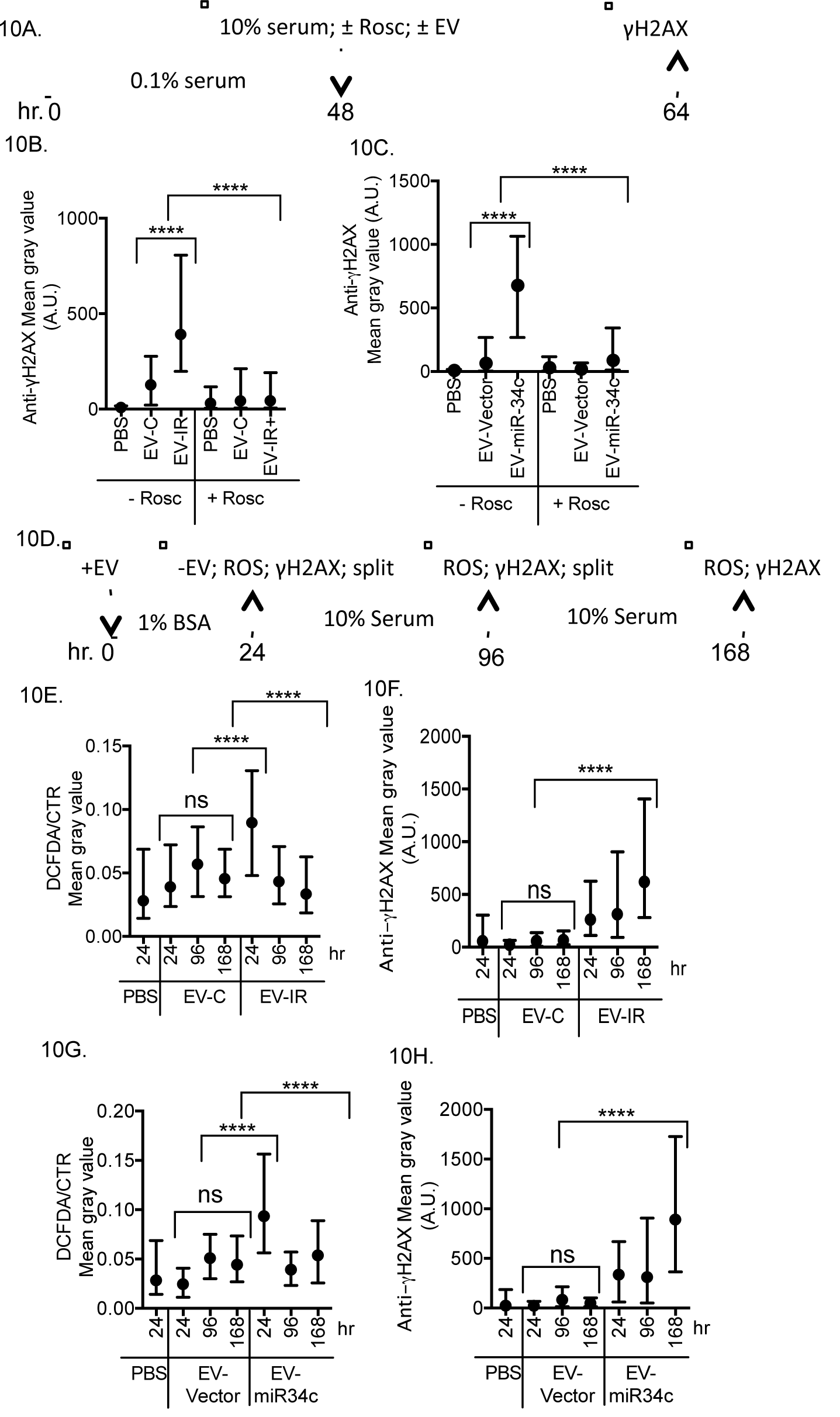
Effects of Roscovitine and Serial Passage on γH2AX Induction by miR-34c-Containing Extracellular Vesicles. (A) Timeline of roscovitine (Rosc) treatment protocol. Responder MEFs were synchronized by serum starvation, released by serum addition with the indicated treatments, and collected for anti-γH2AX staining 16 hours later. (B, C) Roscovitine inhibited the induction of γH2AX by miR-34c-containing EV: Values shown are medians with interquartile ranges of anti-γH2AX mean gray values from at least 200 cells per sample per experiment. \*\*\*\**P*≤ 0.0001. Kruskal-Wallis test. Representative images and anti-γH2AX mean gray values in individual cells are shown in Fig. S5A, B & G, H. (D) Timeline of serial passage protocol. (E, G) ROS increase was lost upon serial passage in the absence of miR-34ccontianing EV: Values shown are medians with interquartile ranges of DCFDA/CTR ratios from at least 200 cells analyzed per sample per experiment. ns: not significant, \*\*\*\**P*≤ 0.0001, Kruskal-Wallis test. Representative images and ratios of individual cells are shown in Fig. S5C, E. (F, H) γH2AX increase was maintained through serial passage in the absence of miR-34c-containing EV: Values shown are medians with interquartile ranges of anti-γH2AX mean gray values from at least 200 cells analyzed per sample. ns: not significant, \*\*\*\**P*≤ 0.0001, Kruskal-Wallis test. Representative images and anti-γH2AX mean gray values of individual cells are shown in Fig. S5D, F & I, J.

### Persistence of EV-IR-Induced γH2AX Increase (Fig. 10; Fig. S5)

An interesting hallmark of bystander DNA damage induced by radiation is the epigenetic propagation of this response (Koturbash et al., 2006). Because EV-IR could increase γH2AX and RAD51 foci in responder cells without causing growth arrest (Fig. 3; Fig. S2), we determined the stability of the ROS and γH2AX responses through serial passages after EV removal (Fig. 10D). The ROS increase detected at the time of EV-IR removal was lost after culturing in full media (Fig. 10E; Fig. S5C). Interestingly, however, the EV-IR-induced increase in γH2AX was stable through two passages in fresh media without EV-IR (Fig. 10F; Fig. S5D, I). The persistence of γH2AX increase was similarly observed when responder cells were treated with EV-miR-34c (Fig. 10 G, H; Fig. S5E, F, J). These results showed that EV-IR-and EV-miR34c-induced increase in γH2AX was a stable response that could be propagated to progeny cells.

## DISCUSSION

### EV from Irradiated Cells Induce Multiple Bystander Effects

This study investigated the role of extracellular vesicles (EV) in radiation-induced bystander effects on colony formation, redox homeostasis and DNA damage. We found that EV isolated from media conditioned by irradiated cells (EV-IR) induced each of those three biological effects in non-irradiated bystander cells but with different efficacies. Although EV-IR was internalized by virtually all responder cells, we found no evidence for cell cycle arrest or senescence in the general population. Rather, EV-IR appeared to only inhibit the colony-forming potential of a small fraction of the responder cells. By contrast, EV-IR did cause ROS levels to rise in virtually all of responder cells. This ROS increase contributed to colony inhibition because an anti-oxidant NAC neutralized the ROS response and reduced the colony-inhibitory activity of EV-IR. However, NAC did not affect EV-IR-induced DNA damage, measured by increases in γH2AX and RAD51 foci and occurring in a sub-population of responder cells. Furthermore, the ROS increase was not maintained after removal of EV-IR whereas the γH2AX increase was propagated to progeny responder cells up to 6 days after EV-IR removal. Together, these results suggest that EV-IR induced multiple biological effects in responder cells by different mechanisms.

### Nuclear Abl Requirement in EV-Mediated Bystander Effects

Results from this study have uncovered a previously unknown function of nuclear Abl in DNA damage response, that is, nuclear Abl is required for irradiated cells to produce miR-34c-containing EV with ROS-and γH2AX-inducing activities. We created the *Abl-µNLS* allele in mice to investigate the physiological role of nuclear Abl in DDR. We show here that IR-induced increase in ROS and DNA damage occurred in the *Abl-µNLS* MEFs despite the lack of nuclear Abl. This may not be a surprising observation since the ROS increase occurs in the cytoplasm through the mitochondria, and the DNA breaks result from physical-chemical reactions. In other studies, we found that nuclear Abl does contribute to the induction of p53-target genes by IR (unpublished), confirming previous results that nuclear Abl activates the p53-family of transcription factors in DNA damage response (Gong et al., 1999; Shaul, 2000). We have also shown that Abl kinase phosphorylates DCGR8 to stimulate precursor miR-34c processing (Tu et al., 2015). In this study, we have identified a previously unknown function of the nuclear Abl-miR34c pathway in EV-mediated bystander effects of radiation. Results shown here establish that irradiated cells require nuclear Abl kinase to include miR-34c in EV for transfer to responder cells. The Abl kinase may passively stimulate miR-34c inclusion as the outcome of its stimulation of miR-34c expression. Alternatively, Abl kinase may actively stimulate miR-34c secretion by targeting this microRNA to micro-vesicles.

### MicroRNA in Radiation-Induced Bystander Effects

Previous studies have found IR to increase the intracellular abundance of many microRNAs by regulating their biogenesis (He et al., 2007a; Mao et al., 2014). Previous studies have also shown that microRNAs affect an array of cellular responses to radiation (Chaudhry, 2014). Results from this study show for the first time that IR-induced miR-34c is secreted in extracellular vesicles to induce ROS and γH2AX in non-irradiated cells. The miR-34-family of microRNAs can target many pro-mitogenic and pro-survival genes to cause growth arrest and apoptosis (He et al., 2007b; Maroof et al., 2014). However, we found that EV-IR and EV-miR-34c did not cause growth arrest or apoptosis in the responder cells, indicating that the previously validated pro-mitogenic and pro-survival mRNAs may not be sufficiently reduced by EV-mediated delivery of miR-34c. Computational analyses have predicted hundreds of other miR-34c targets that could be involved in the observed induction of ROS or γH2AX. Furthermore, miR-34c may trigger a cascade of gene expression alterations or it may collaborate with other EV-delivered factors to increase ROS and γH2AX in responder cells. Our findings that nuclear Abl is essential for irradiated cells to produce EV-IR with ROS-and γH2AX-inducing activities, but the miR34-family is essential only for the ROS-inducing activity suggest that EV-IR must contain other nuclear-Abl-dependent DNA damage-inducers that remain to be identified.

## MATERIALS & METHODS

### Cell Lines

Fibroblasts were derived from *Abl^+/+^* (*Abl-wt*) or littermate *Abl^µ/µ^*(*Abl-µNLS)* mouse embryos. The *Abl-µNLS* allele was generated by knock-in mutations to substitute the eleven lysines and arginines in the three nuclear localization signals (NLS) with glutamine (Preyer et al., 2007). The *Abl-wt* and *Abl-µNLS* mouse embryo fibroblasts (MEFs) were immortalized by serial passages, and these MEFs do not express p53. Primary, non-immortalized, MEFs from *miR-34a, b, c-*triple knockout mice (miR34^TKO^) and wild-type littermates (miR34^WT^) (Concepcion et al., 2012) were irradiated between passages 3 and 6. MEFs and HEK293T cells (Thermo Fisher Scientific) were cultured in DMEM high glucose media with 10% fetal bovine serum (FBS) and antibiotics.

### Irradiation

Cells were exposed to 10 Gy of gamma-irradiation using Mark I model 50 irradiator with Cesium 137 isotope as source (Maker: J.L. Shepherd & Associates).

### Isolation of Extracellular Vesicles (EV)

To avoid EV from fetal bovine serum (FBS), 10^7^ cells (in ten 10-cm dishes) were switched to FBS-free media with 1% BSA two hours before irradiation. At 24 hours after irradiation, the media were collected for EV isolation by differential ultracentrifugation as previously described (Thery et al., 2006) (Fig. 1A). The pelleted EV fraction was washed and re-suspended in 300 µl of phosphate buffered saline (PBS) and stored in 50 µl aliquots at −80°C. For isolation of EV from HEK293T cells, supernatant-1 collected after the 2000xg spin (Fig. 1A) was filtered through a 0.45-micron filter (Corning) before continuing onto the next steps of ultracentrifugation. Protein content of EV was determined by the Lowry method.

### Nanoparticle Tracking Analysis

Nanosight LM-10HS was used for nanoparticle tracking analysis. This analysis uses the diffraction measurement of Brownian motion of particles. The EV suspension was diluted 300 fold in PBS and 1 µl of the diluted suspension was video taped by Nanosight to determine the size distribution and the concentration of particles. Each EV preparation was analyzed in triplicates as previously described (Akers et al., 2016).

### Uptake of Extracellular Vesicles

EV suspensions were incubated with PKH26, a fluorescent membrane-binding dye (Sigma Aldrich, St. Louis) for 5 min at room temperature, followed by addition of 1% BSA, and then centrifuged at 100,000×g for 70 min to isolate PKH26-labeled EV as previously described (Mineo et al., 2012). Responder MEFs were incubated with PKH26 in PBS, or PKH26-labeled EV-C or PKH26-labeled EV-IR (25 µg each). After 3 or 24 hours, cells were fixed with 4% para-formaldehyde (PFA) for 20 min at room temperature and counterstained with Hoechst 33342. Cells were viewed using an Olympus FV1000 Spectral Confocal microscope. No fluorescence was detected in cells incubated with PKH in PBS. Using FIJI (ImageJ), we measured the PKH26 mean gray values from at least 200 cells per treatment and calculated the mean and standard deviations. The number of PKH26-positive cells was counted by eye, and percentages were calculated from PKH26-positive cells over total number of nuclei.

### Colony Formation Assay

Responder cells (*Abl-wt* MEFs) were seeded at 1000 cells per 6-cm plate. Media was changed to 1% BSA without FBS before incubation with EV. After 24 hours, cells were switched back to media with 10% FBS and cultured for 15 days with media refreshed every other day. The colonies were fixed with 100% methanol and stained with 0.05% crystal violet. Excess dye was removed & plates were left to dry over-night. Cluster of more than 50 cells were considered as colonies. Survival fraction was calculated as colonies/cells seeded with the survival fraction in PBS treated plates set to 1. Images of the colonies were acquired using Alpha imager HP System.

### Reactive Oxygen Species (ROS) Assay

ROS was measured using the ROS-ID kit (Enzo Life Sciences, Farmingdale) according to manufacturer’s protocol. Live cells were also stained with Cell Tracker Red (CTR) (Molecular Probe) as a control for cell volume. Responder cells were seeded into chamber slides, incubated with EV in serum-free media containing 1% BSA for 24 hours, then stained with CTR and DCFDA. Immediately after dye addition, live cell images were captured using an Olympus FV1000 Spectral Confocal Microscope for CTR (Channel 3) & DCFDA (Channel 1). FIJI (ImageJ) software was used to create masks from channel 3 (CTR), and then the masks were transferred onto channel 1 (DCFDA). The mean gray values (MGVs) in channels 1 and 3 were recorded within the masks, and the DCFDA/CTR MGV ratio was calculated for each mask. See Figs. S3D; S4C, G for plots of ranked DCFDA/CTR ratio of individual cell from representative experiments. From each experiment, we collected the ratio from at least 200 cells per sample. We then determined the median and the interquartile range of ratios collected from one to three experiments (200-600 cells) as indicated in the figure legends.

### Immunofluorescence

Acid-washed coverslips stored in 100% ethanol were placed in 24-well plates and approximately 20,000 MEFs were seeded per well. After incubation with EV for 24 hours in serum-free media containing 1% BSA, cells were fixed in 4% PFA for 15 min, washed with 0.02% Tween-20 in Tri-buffered saline (TBS) twice (5 min each), permeablized with 1% Triton X-100 in TBS for 15 min and then blocked with 5% BSA for 30 min at room temperature. The coverslips were incubated with primary antibody for 1 hour in 37°C: anti-Abl (8E9) (6µg/ml) from ThermoFisher Scientific, anti-phospho-Ser139-H2AX (anti-γH2AX, 1/400) from Cell Signaling, anti-RAD51 (1/50) from Santa Cruz. Coverslips were washed twice with 0.02% Tween-20 in TBS twice (5 min each) and then incubated with ALEXA fluor-488 (Invitrogen)-chicken anti-mouse (1/500) or ALEXA fluor-594 (Invitrogen)-donkey anti-rabbit (1/500) at 37°C for 30 min. Nuclei were stained with Hoechst 33342. Coverslips were mounted with Prolong Gold Antifade Reagent and sealed with nail polish before imaging. Images were captured using an Olympus FV1000 Spectral Confocal Microscope.

### Quantification of γH2AX

FIJI (ImageJ) software was used to create masks from channel 0 (Hoechst 33342), the masks were then transferred onto channel 1 (anti-γH2AX), and the channel 1 mean gray value (MGV) within each mask was recorded. See Figs. S3B, C; S4D, H; S5G, H, I, J for plots of ranked γH2AX MGV of individual cell from representative experiments. From each experiment, we collected the MGV from at least 200 cells per sample. We then determined the median and the interquartile range of MGV collected from one to three experiments (200-600 cells) as indicated in the figure legends.

### Senescence-Associated β-Galactosidase Staining

Irradiated (10 Gy) or EV-treated (25µg, 24 hours) MEFs were cultured for additional 5 days in serum-supplemented media and counterstained with Hoechst 33342 and the Senescence Cells Histochemical Staining Kit (Sigma-Aldrich). The number of nuclei was counted in images captured by the KEYENCE BZ-X700 All-in-One Fluorescence Microscope at a magnification of 20X, with 50% transmitted light, and white balance red-blue-green areas of 1.46, 0.98, and 1.24, respectively. The β-galactosidase positive blue cells were counted under Phase contrast microscope. At least 200 cells were scored per sample per experiment to calculate the percentage of positive senescent cells.

### Immunoblotting

Cell pellets were lysed in RIPA buffer (25mM Tris-HCl pH 7.6, 10% Glycerol, 1% NP40, 0.5% sodium deoxycholate, 1x Protease inhibitors (Roche), 150mM Sodium chloride, 50mM Sodium fluoride, 10mM Sodium beta-glycerophosphate, 10mM Sodium orthovandate, 10mM sodium pyrophosphate, 1mM PMSF). Proteins were separated using SDS-PAGE & transferred onto Nitrocellulose membranes (Millipore). Membranes were blocked for 1 hour at room temperature, incubated with anti-Abl (8E9) (1/500) & anti-actin (1/2000) from Sigma Aldrich for 1 hour, washed and incubated with secondary antibody (Anti-mouse: HRP-linked) & developed using ECL reagents (Pierce).

### Cell Cycle Analysis

Cells were collected at 24 hours post irradiation or EV treatment by trypsinization, followed by centrifugation at 1400 RPM for 6 min. Cell-pellets were fixed in ice cold 70% ethanol overnight and stained with 40 µg/ml propidium iodide (PI) (Sigma) and 100 µg/ml RNaseA (Sigma) for 30 min at 37^o^C in the dark. PI staining was analyzed using Sony SH800 FACS sorter and software.

### Retrovirus Packaging and Infection

Abl^WT^ & Abl^µNLS^ were stably expressed in *Abl-µNLS* MEFs by retroviral infection. BOSC23 cells were transfected with retroviral vector pMSCV expressing Abl^WT^ or Abl^µNLS^. Culture media collected at 48 hours after transfection was filtered and added to *Abl-µNLS* MEFs with polybrene (4µg/ml). Infected cells were then selected for resistance to hygromycin (150 µg/ml).

### Transfection

Genetran (Biomiga) was used to transfect HEK239T cells with miR-34cminigene and pCDNA3-AblPPn plasmid DNA (Tu et al., 2015). Transfected cells and their media (for EV isolation) were collected 24 hours after transfections.

### RNA measurements

The SeraMir Exosome RNA amplification kit (System Biosciences) was used to extract RNA from EV pellets. Total cellular RNA was extracted using TRIzol (Life Technologies). Synthesis of cDNA was carried out using ABI reverse Transcription kit (Life Technologies). For measurements of mature miR-34c, stem-loop primer was used for reverse transcription (Tu et al., 2015). U6 was used as the reference gene for normalization of miR-34c abundance. GAPDH was used as reference gene for normalization of p21Cip1, Pdgfrb, Pdgfra & Myc abundance. Realtime PCR reactions were carried out using StepOnePlus system. Subtraction of the reference gene CT value from the experimental gene CT value generated the normalized ΔCT. Relative abundance was then calculated as 2-^ΔΔ^CT, where ΔΔCT values were ΔCT of sample subtracted by ΔCT of vehicle-treated or vector transfected cells. Primer sequences: U6-F: CTCGCTTCGGCAGCACA, U6-R: AACGCTTCACGAATTTGCGT, Stem-loop miR34c: GTCGTATCCAGTGCAGGGTCCGAGGTATTCGCACTGGATACGACGCAATC, q-miR34c-F: AGGCAGTGTAGTTAGCTG, q-miR-R: GTGCAGGGTCCGAGGT, p21-F: CCATGTGGACCTGTCACTGTCTT, p21-R: AGAAATCTGTCATGCTGGTCT, Pdgfrb F: GTTGTTGCTGTCCGTGTTATG, Pdgfrb R: GGCCCTAGTGAGTTGTTGTAG, Myc F: CGACTCTGAAGAAGAGCAAGAA, Myc R: AGCCAAGGTTGTGAGGTTAG, Pdgfra F: CTCAGAGAGAATCGGCCCCA, Pdgfra R: CACCAGCCTCCCGTTATTGT

### Statistical Analysis

The statistical analyses were performed using Graph-Pad Prism 6. For clonogenic survival & qRT-PCR measurements, the mean ± SD from three independent experiments were analyzed using ONE WAY ANOVA. For ROS and γH2AX measurements, the ratios(ROS)or the mean gray values (γH2AX)from 200-600 cells from one to three independent experiments per sample were ranked across samples and the mean ranks analyzed using the non-parametric Kruskal-Wallis test. For each statistical test, ns: not significant; \**P≤*0.05, \*\**P≤*0.01, *** *P≤*0.001, **** *P≤*0.0001.

## SUMMARY

Ionizing radiation stimulates nuclear accumulation of Abl tyrosine kinase that is required for directly irradiated cells to produce microRNA-34c-containing extracellular vesicles which transfer the microRNA into non-irradiated cells to induce reactive oxygen species and bystander DNA damage.

## ACKNOWLEDGEMENTS

We thank Chenhui Bian, Neal Shah and Louis Nguyen for excellent technical assistance. We thank Dr. Andrea Ventura, Memorial Sloan Kettering Cancer Centre, New York for the miR34^WT^ and miR34^TKO^ primary MEFs. We thank Dr. Clarke Chen & Dr. Johnny Akers for helping with the Nanoparticle Tracking Analysis.

## AUTHOR CONTRIBUTIONS

SR designed and performed the experiments, analyzed the data, and wrote the paper. A.H, J.C, performed experiments and analyzed the data. JYJW conceived of the idea,designed the experiments, analyzed the data and wrote the paper.

## SUPPLEMENTAL MATERIAL

### Nuclear Abl Drives miR-34c Transfer by Extracellular Vesicles to Induce Radiation Bystander Effects

*S. Rastogi et al*

**Figure S1.** Representative Images of Clonogenic Assay Results (supporting Figs. 2 & 3).

**Figure S2.** Extracellular Vesicles from Irradiated Cells Did Not Induce p21Cip1, Cell Cycle Arrest or Senescence (supporting Fig. 2).

**Figure S3.** Treatment with EV-IR but not EV-C Induced ROS, γH2AX, and RAD51 Foci in Responder Cells (supporting Fig. 3).

**Figure S4.** ROS- and γH2AX-Inducing Activities of EV Preparations from Transfected HEK293T cells (supporting Fig 9).

**Figure S5.** Effects of Roscovitine and Cell Passage on γH2AX (supporting Fig 10).

**Figure S1.**
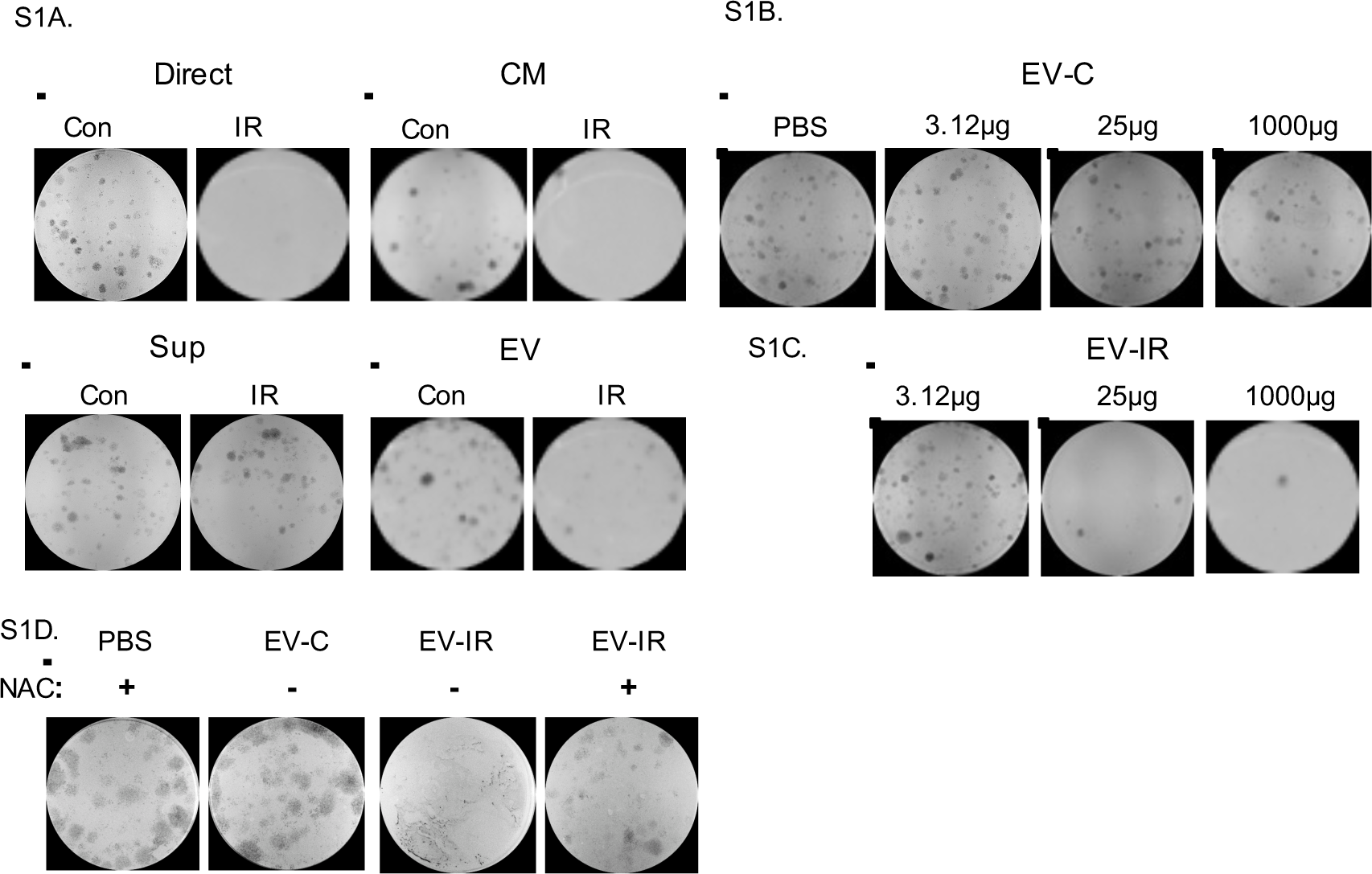
Representative Images of Clonogenic Assay Results (supporting Figs. 2 & 3). (A) Representative images of colonies for the data shown in Fig 2A. Direct: direct irradiation. CM: conditioned media. Sup: supernatant-2 from CM (see Fig. 1A). EV: washed extracellular vesicle pellet from CM (see Fig. 1A) Con: control, no irradiation. IR: ionizing radiation, 10 Gy. (B) Representative images of colonies for the data shown in Fig 2B. EV-C: EV from CM of non-irradiated MEFs. (C) Representative images of colonies for the data shown in Fig 2C. EV-IR: EV from CM of irradiated MEFs. (D) Representative images of colonies for the data shown in Fig 3C. NAC: N-acetyl-cysteine (5 mM)

**Figure S2.**
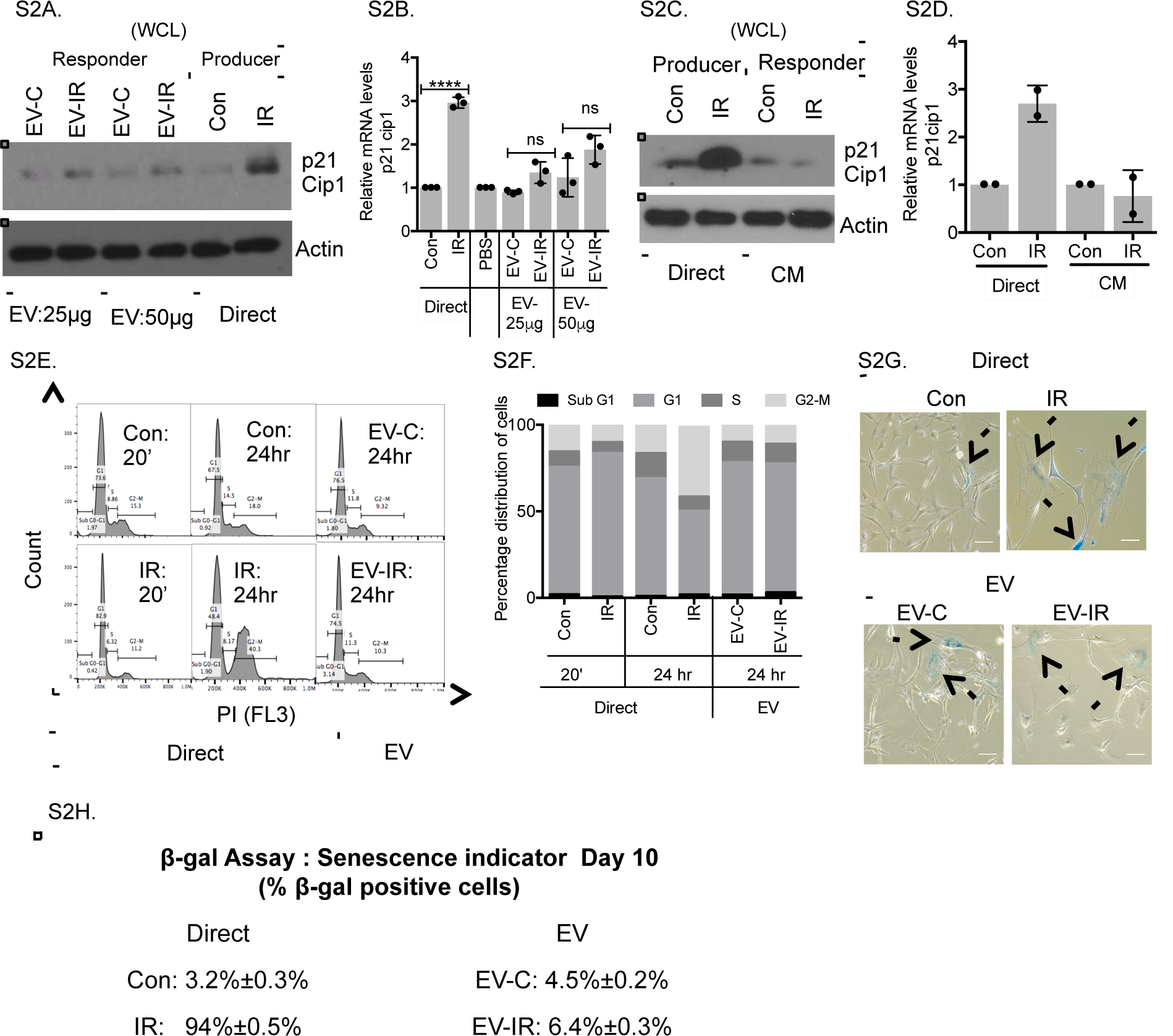
Extracellular Vesicles from Irradiated Cells Did Not Induce p21Cip1, Cell Cycle Arrest or Senescence (supporting Fig. 2). (A) Western blotting with anti-p21Cip1 or anti-actin (loading control) of whole cell lysates (WCL) from non-irradiated responder MEFs after 24 hours of treatment with the indicated EV at two different concentrations (25 μg & 50 μg) or of WCL from non-irradiated (Con) or irradiated (IR, 10 Gy) MEFs that produced the EV at 24 hours after direct irradiation. Note that EV-IR did not induce p21Cip1 protein in non-irradiated responder MEFs. (B) Relative abundance of p21Cip1 RNA in non-irradiated (Con) or directly irradiated (IR, 10 Gy) MEFs at 24 hours, or in non-irradiated responder MEFs at 24 hours after treatment with PBS or the indicated EV at two different concentrations (25 μg & 50 μg). Data shown are mean ± SD of relative abundance from three independent experiments with the normalized p21Cip1 levels in Con or PBS-treated MEFs set to 1. ns: not significant, **** *P*< 0.0001, ONE WAY ANOVA. (C) Western blotting with anti-p21Cip1 or anti-actin (loading control) of whole cell lysates (WCL) from MEFs at 24 hours after direct irradiation or not (producer) or after treatment with CM from control or irradiated cells (responder). Note that CM-IR did not induce p21Cip1 protein in nonirradiated responder MEFs. (D) Relative abundance of p21Cip1 RNA in MEFs as treated in (C). Data shown are mean ± SD from two independent experiments with p21Cip1 levels in non-irradiated cells or non-irradiated- CM-treated cells set to 1. (E) FACS analysis of DNA content: MEFs were collected at 20 minutes or 24 hours after 10 Gy irradiation (IR) or not (Con), or at 24 hours after treatment with EV-C or EV-IR (25 μg each). Representative FACS histogram plots with cell cycle distribution determined by the software FlowJo are shown. (F) Stacked bar graph summarizing the cell cycle distribution of MEF populations after the indicated treatments from one representative experiment. The immortalized *Abl-wt* MEFs are p53-deficient and did not undergo G1 arrest. Direct irradiation but not EV-IR induced G2 arrest. (G) Directly irradiated or EV-treated (25 μg, 24 hours) MEFs were cultured for 10 days and stained for β-galactosidase. Representative images with β-galactosidase-positive (blue) cells are marked with arrows (Scale bar: 30 μm). (H) Quantification of β-galactosidase-positive cells among 200 cells scored per sample per experiment. Values shown in are mean ± SD from two independent experiments. Direct irradiation with 10 Gy IR caused >90% of Abl-wt MEFs to become β-galactosidase-positive, but treatment with EV-C or EV-IR did not induce senescence.

**Figure S3.**
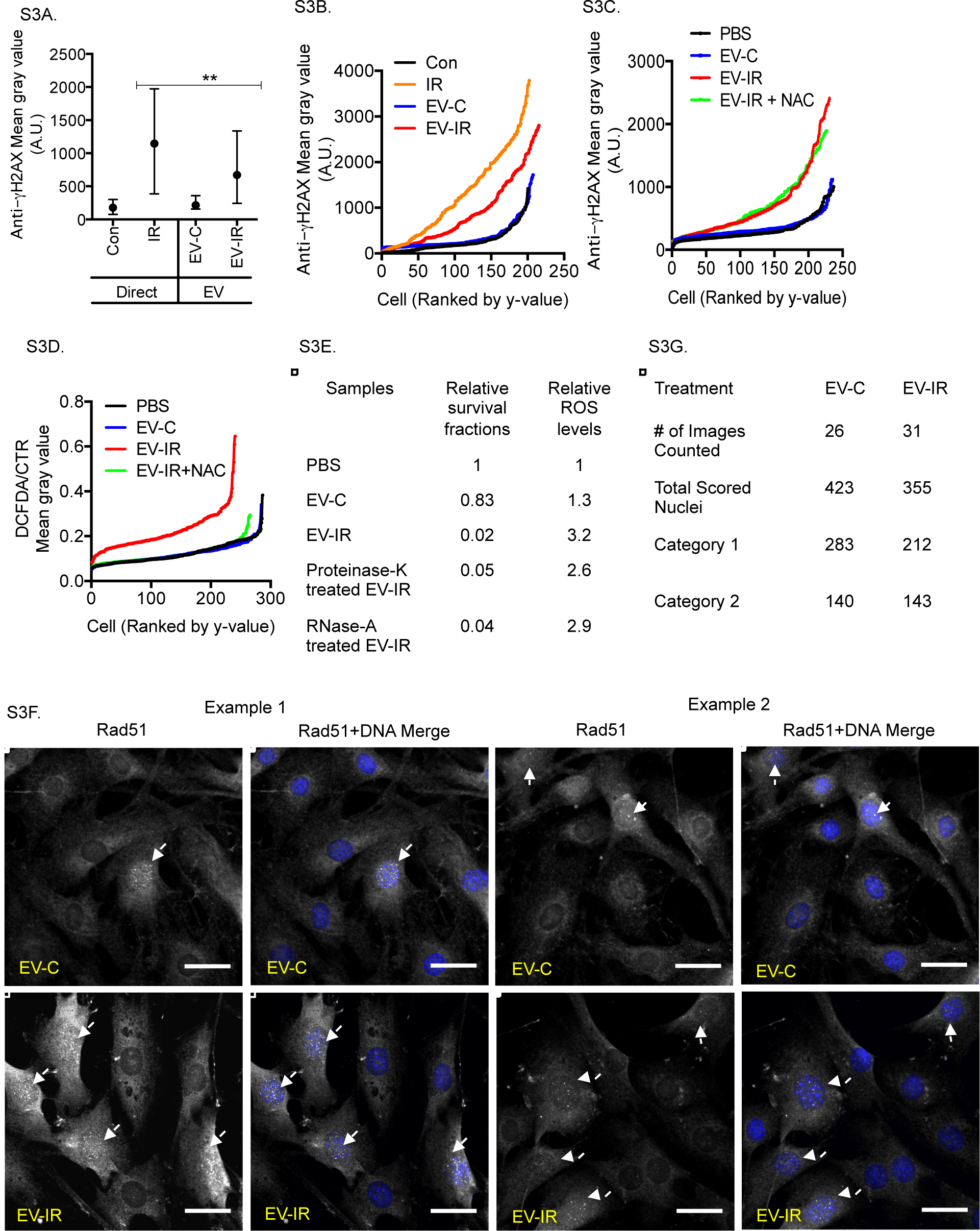
Treatment with EV-IR but not EV-C Induced ROS, γH2AX, and RAD51 Foci in Responder Cells (supporting Fig. 3). (A) Induction of γH2AX by direct irradiation (10 Gy) or by treatment with EV-IR (3.7 μg) : Values shown are medians with interquartile ranges of anti-μH2AX mean gray values from 200 cells analyzed per sample. **P. 0.01. Kruskal-Wallis test. (B) The γ-H2AX mean gray values of individual cells for the data shown in (A). (C) The γ-H2AX mean gray values of individual cells from one of the three experiments shown in Fig. 3D. (D) The ratio of DCFDA/CTR mean gray values of individual cells from one of the three experiments shown in Fig. 3A. (E) Proteinase K and RNaseA treatment did not inactivate the colony-inhibitory or the ROS-inducing activities of EV-IR. Relative survival fractions and median ROS levels in responder MEFs after the indicated treatments for 24 hours. Isolated EV were incubated with proteinase K (0.05 mg/ml) for 10 minutes at 60°C or with RNaseA (0.5 mg/ml) for 20 minutes at 37°C to degrade unprotected protein & RNA. (F) Non-irradiated responder MEFs treated with EV-C or EV-IR (25 μg each) for 24 hours were fixed and stained with anti-RAD51 (gray) and Hoechst 33324 (DNA; blue). Two representative images are shown for each treatment. The white arrows point to nuclei with RAD51 foci (Scale bar: 35μm). (G) Summary of total nuclei scored for the data shown in Fig. 3F.

**Figure S4.**
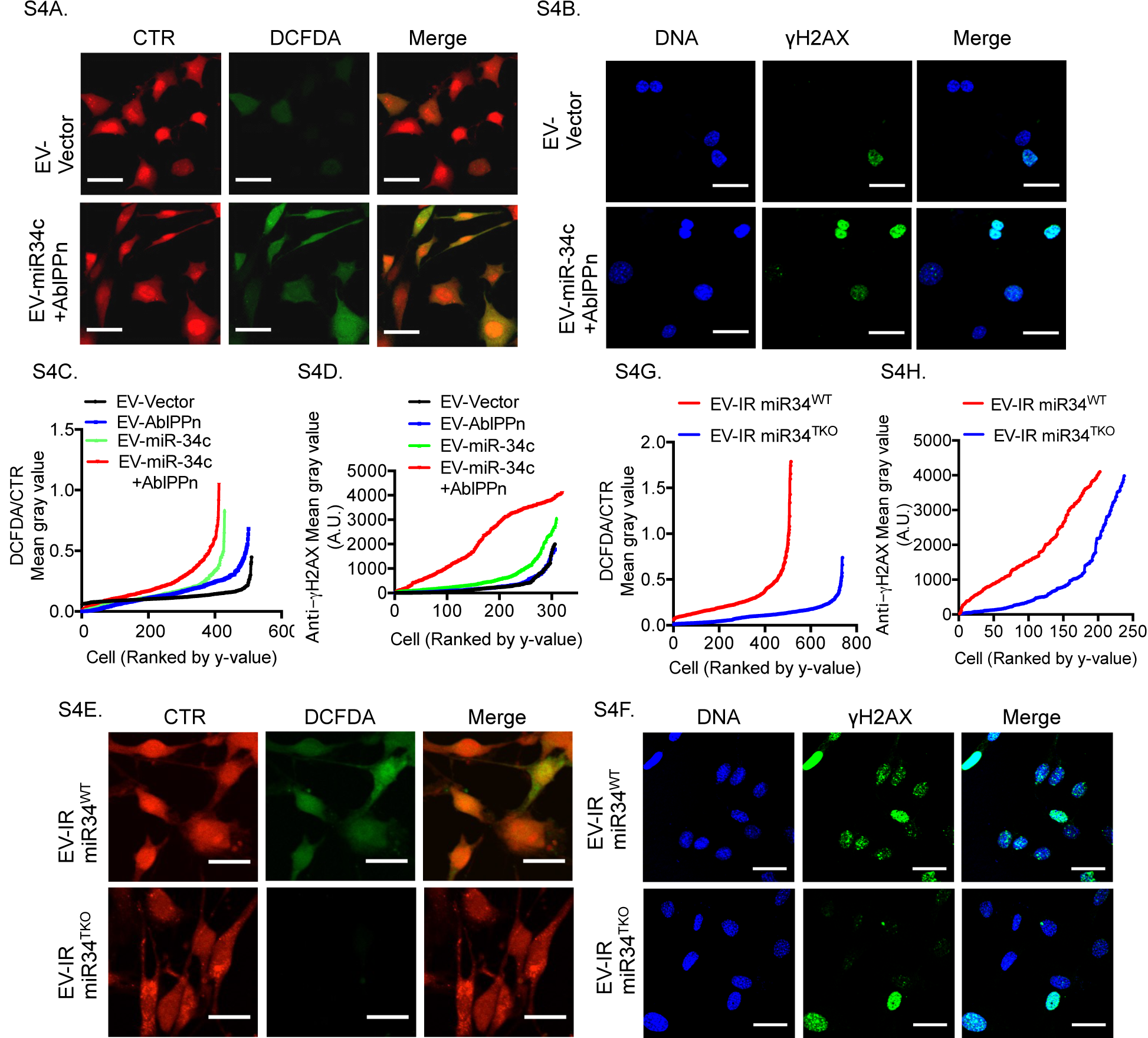
ROS- and γH2AX-Inducing Activities of EV Preparations from Transfected HEK293T cells (supporting Fig 9). (A, C) Representative images (A)(Scale bar: 35 μm) and DCFDA(green)/CTR(red) ratios of individual live cells (C) at 24 hours after treatment with the indicated EV (25 μg each). The medians and interquartile ranges of the DCFDA/CTR ratios are shown in Fig. 9F. (B, D) Representative images (B)(Hoechst 33324, DNA: blue; anti-γH2AX: green; Scale bar: 35 μm) and anti-γH2AX mean gray values of individual cells (D) after 24 hours of treatment with the indicated EV (25 μg each). The medians and interquartile ranges of the anti-γH2AX mean gray values are shown in Fig. 9G. (E, G) Representative images (E) (Scale bar: 35 γm) and DCFDA(green)/CTR(red) ratios of individual live cells (G) after 24 hours of treatment with EV-IR (25 γg each) isolated from media conditioned by irradiated miR34^WT^ or miR34^TKO^ MEFs. The medians and interquartile ranges of DCFDA/CTR ratios are shown in Fig. 9H. (F, H) Representative images (F) (Hoechst 33324, DNA: blue; anti-γH2AX: green; Scale bar: 35 μm) and anti-γH2AX mean gray values of individual cells (H) after 24 hours of treatment with the indicated EV-IR (25 μg each) isolated from media conditioned by irradiated miR34^WT^ or miR34^TKO^ MEFs. The medians and interquartile ranges of anti-γH2AX mean gray values are shown in Fig. 9I.

**Figure S5.**
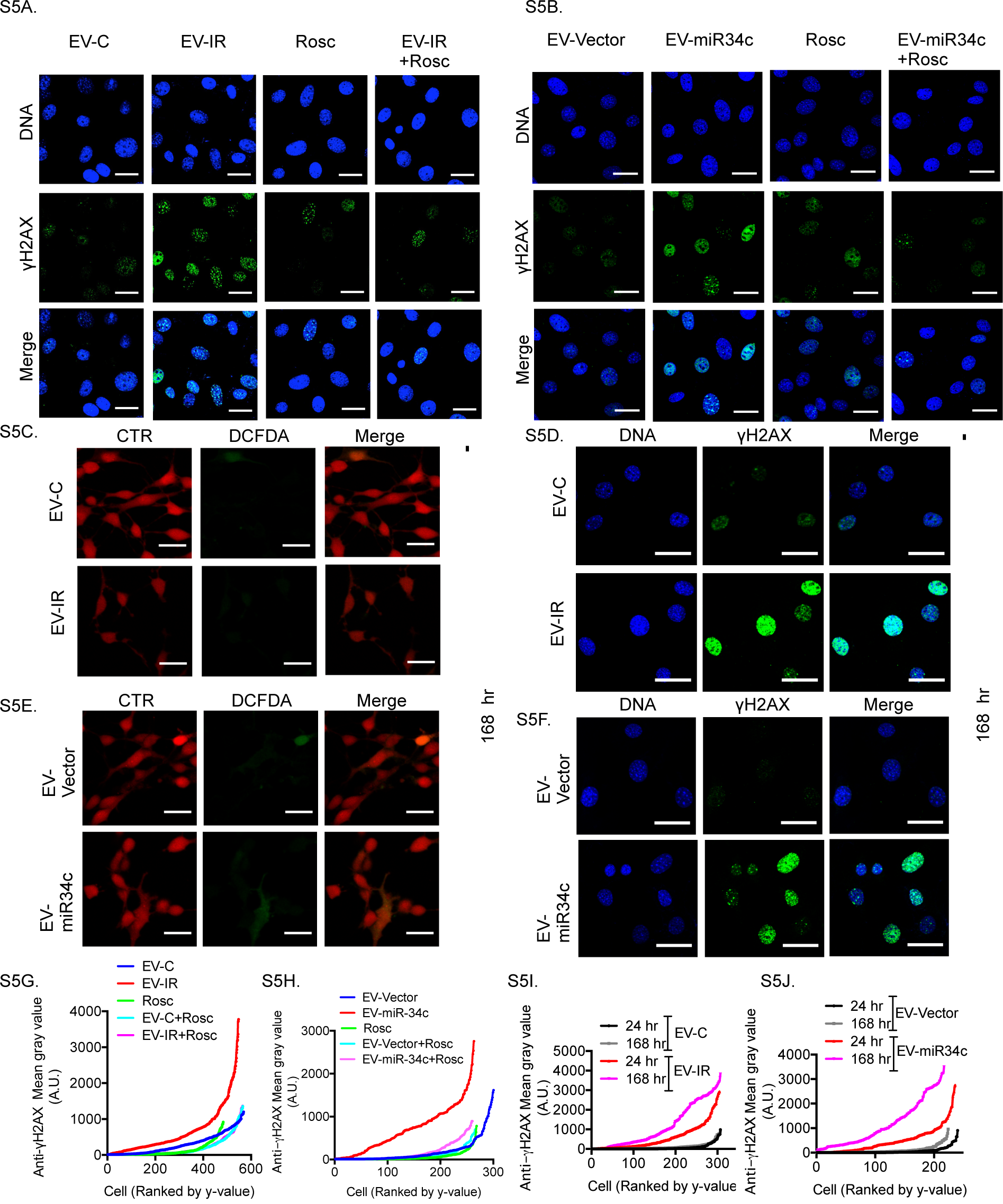
Effects of Roscovitine and Cell Passage on γH2AX (supporting Fig 10). (A, G) Representative images (A) (Hoechst 33324, DNA: blue; anti-γH2AX: green; Scale bar: 35 μm) and anti-γH2AX mean gray values in individual cells (G) for data shown in Fig 10B. (B, H) Representative images (B) (Hoechst 33324, DNA: blue; anti-γH2AX: green; Scale bar: 35 μm) and anti-γH2AX mean gray values in individual cells (H) for data shown in Fig 10C. (C, E) Representative images of live responder cells stained with CTR (red), and DCFDA (green) at 168 hours after treatment with (C) EV-C or EV-IR from control or irradiated (10 Gy IR) Abl-wt MEFs, or (E) EV-vector or EV-miR-34c from vector or miR-34c-minigene transfected HEK293T cells (Scale bar: 35μm). The medians and interquartile ranges of DCFDA/CTR ratios are shown in Fig. 10E,G. (D, I) Representative images (D) (Hoechst 33324, DNA: blue; anti-γH2AX: green, Scale bar: 35 μm) and anti-γH2AX mean gray values of individual cells (I) at 168 hours after treatment with EV-C or EV-IR The medians and interquartile ranges of anti-γH2AX mean gray values are shown Fig 10F. (F, J) Representative images (F) (Hoechst 33324, DNA: blue; anti-γH2AX: green, Scale bar: 35 μm) and anti-γH2AX mean gray values of individual cells (J) at 168 hours after treatment with EV-vector or EV-miR34c. The medians and interquartile ranges of anti-γH2AX mean gray values are shown Fig 10H.

